# A new vulnerability to BET inhibition due to enhanced autophagy in BRCA2 deficient pancreatic cancer

**DOI:** 10.1101/2023.05.30.542934

**Authors:** EunJung Lee, Suyakarn Archasappawat, Keely Ji, Jocelyn Pena, Virneliz Fernandez-Vega, Ritika Gangaraju, Nitin Sai Beesabathuni, Martin Jean Kim, Qi Tian, Priya Shah, Louis Scampavia, Timothy Spicer, Chang-Il Hwang

## Abstract

Pancreatic cancer is one of the deadliest diseases in human malignancies. Among total pancreatic cancer patients, ∼10% of patients are categorized as familial pancreatic cancer (FPC) patients, carrying germline mutations of the genes involved in DNA repair pathways (*e.g., BRCA2*). Personalized medicine approaches tailored toward patients’ mutations would improve patients’ outcome. To identify novel vulnerabilities of *BRCA2*-deficient pancreatic cancer, we generated isogenic *Brca2*-deficient murine pancreatic cancer cell lines and performed high-throughput drug screens. High-throughput drug screening revealed that *Brca2*-deficient cells are sensitive to Bromodomain and Extraterminal Motif (BET) inhibitors, suggesting that BET inhibition might be a potential therapeutic approach. We found that *BRCA2* deficiency increased autophagic flux, which was further enhanced by BET inhibition in *Brca2*-deficient pancreatic cancer cells, resulting in autophagy-dependent cell death. Our data suggests that BET inhibition can be a novel therapeutic strategy for *BRCA2*-deficient pancreatic cancer.

## Introduction

Pancreatic cancer is the 3^rd^ leading cause of cancer-related deaths in the United States, with a dismal 5-year survival rate of 11% (1). The low survival rate of pancreatic ductal adenocarcinoma (PDAC) is largely attributed to the late diagnosis of the disease when cancer has metastasized, making surgical resection a viable option for less than 20% of PDAC patients (2). In addition, the highly resistant and heterogeneous nature of PDAC makes the benefits from the first-line chemotherapies rather modest. Therefore, there is an urgent need to develop more effective therapies for patients with PDAC.

PDAC is driven by somatic mutations in the oncogene *KRAS* and the tumor suppressor *TP53* (3). In addition, around 10% of PDAC cases are hereditary and are categorized as familial pancreatic cancer (FPC), which describes families with two or more first-degree relatives afflicted by PDAC (4). Previously, sequencing of FPC patient samples has revealed that FPC patients harbor germline mutations in genes related to DNA repair pathways (hereinafter referred to as ‘FPC genes’) which increase the likelihood of developing PDAC in their lifetime (4). The most widely studied FPC gene is *BRCA1/2*, which code for key proteins mediating the homologous recombination (HR) DNA repair pathway (5). Defects in BRCA1/2 create a unique set of vulnerabilities, such as increased genomic instability and defects in the HR pathway, which augments sensitivity to platinum-based chemotherapies and increases the reliance on alternative DNA repair pathways, respectively (6). In the past decades, studies have found that these unique vulnerabilities inherent in *BRCA1/2* mutant cancers (e.g., breast, ovarian, and pancreatic cancers) could be utilized to induce synthetic lethality (7). Other FPC gene mutations which can cause a defect in HR pathway (termed ‘BRCAness’) are thought to get benefits from similar approaches. Indeed, Pancreas Cancer Olaparib Ongoing (POLO) clinical trial has found that Poly ADP-ribose Polymerase inhibitor (PARPi) showed significant benefit in *BRCA1/2* mutant PDAC patients (8). The success of PARPi as a targeted therapy for *BRCA1/2* mutant PDAC patients highlighted the need to develop other forms of targeted therapies that could exploit the unique vulnerabilities within FPC patients. This, in turn, will allow for the treatment of a wider range of patients with a larger range of FPC mutations.

To identify novel targeted therapies that could induce synthetic lethality in *BRCA2*-deficient PDAC, we established isogenic *BRCA2*-deficient murine PDAC cell lines, performed a high-throughput screening (HTS) of drugs, and validated the drug responses. From this screening, we identified that *BRCA2* deficiency confers increased sensitivity to Bromodomain and Extra terminal Motif Domain (BET) inhibitors (BETi). These BET proteins contain two tandem bromodomains in N-terminal, which can bind to lysine-acetylated histones and regulate gene transcription (9). Further genetic and pharmacological perturbation study revealed that BET inhibition induces increased cell death in *BRCA2*-deficient PDAC, through promoting autophagy-dependent cell death.

## Materials and Methods

### Cell Lines and Cell Culture Condition

Murine KPC cell lines (KPC-mT3 and KPC-mT19) were maintained with DMEM (Corning, 10-013-CV) supplemented with 10% FBS (Gen Clone, 25-550H), 1% penicillin-streptomycin (P/S, Gibco, 15140-122) (10). Human pancreatic cancer cell lines (CAPAN1, PANC1, CFPAC1, CAPAN2, and MiaPaCa-2) were obtained from ATCC. All human cell lines were cultured in RPMI 1640 (Corning, 10-040-CV) supplemented with 10% FBS and 1% P/S.

### Compound Library and HTS

A collection of ∼3,300 clinically approved drugs obtained from multiple vendors were assembled at the UF Scripps Biomedical Research High-throughput Screening Center and reformatted into 1536-well source plates for automated robotics screening (11). In addition, the NCI-approved oncology drug set of 133 compounds was obtained directly from the NCI, described previously (12, 13).

### Cell Viability Assay and Cytotoxicity Assay

1500 cells (for mouse KPC cells) or 3000 cells (for human PDAC cell lines) in 50 µL of complete DMEM were plated per well of 96 well plates. 24 hrs post-plating, corresponding drugs were serially diluted in complete DMEM. 50 µL of diluted drugs were treated to cells and cells were incubated. 72 hrs post drug treatment, media was carefully removed. Then, 100 µL of diluted AlamarBlue solution prepared with 180 µL of Resazurin (Fisher Scientific, AC418900010) in 50 mL of 1x PBS was carefully added to each well. Cells with AlamarBlue solution were incubated in the 37°C 5% CO_2_ incubator for 2 hrs. Absorbance at 570 nm and 600 nm was measured with a plate reader. Using the PRISM software, the cell viability measurements from each cell line were normalized to measurement from vehicle treated wells, then cell viability curves were generated using nonlinear regression (curve fit). Drug combination effect was analyzed using the Combenefit software (14). To measure the cytotoxicity, the CytoTox-Glo assay (Promega, G9291) was used, and normalized by the cell viability based on CellTiter-Glo Luminescent cell viability assay. Luminescence signal was read by the SpectraMax iD5 plate reader (Molecular Device).

### RNA-seq and GSEA

For RNA-seq with the FPC gene KO cells and control cells, cells were plated, then treated with 1 µM of JQ1 or DMSO for 72 hrs. Cells were lysed with 1 mL of TRIzol (Thermo Fisher Scientific, 15596026) and 200 µL of Chloroform following the manufacturer’s instructions. The collected RNA was purified with PureLink RNA Mini Kit (Invitrogen, 12183018A) following the manufacturer’s protocol. RNA concentration was determined with nanodrop. Library preparation and RNA-sequencing were performed by Novogene Co., LTD (Beijing, China). In brief, mRNA was enriched using oligo(dT) beads, and rRNA was removed using the Ribo-Zero kit. The mRNA was fragmented, and cDNA was synthesized by using an mRNA template and random hexamers primer, after which a second-strand synthesis buffer (Illumina), dNTPs, RNase H, and DNA polymerase I were added for the second-strand synthesis, followed by adaptor ligation and size selection. The library was sequenced by the Illumina Novaseq platform. Raw data were aligned to mm10 genome using HISAT2, read counts and normalized read counts were generated using the featureCounts, and the differentially expressed genes were identified using DESeq2. Principal Component Analysis plots were generated utilizing packages “factoMineR” and “factoextra” in R utilizing normalized counts of RNA-seq. Gene set enrichment analysis was performed with 7658 gene sets contained in C5: Gene Ontology Biological Processes database with normalized RNA-seq data.

### Western Blotting

Cells were trypsinized into single cells to yield 10^6^ cells. To prepare whole cell lysate, cell pellets were lysed with 1% TNET buffer (50 mM Tris pH 7.5, 1% Triton X-100, 150 mM NaCl, 5 mM EDTA) with 1× protease inhibitor cocktails (Sigma-Aldrich, Sial-11836170001), and phosphatase inhibitor cocktails (Roche, 04906837001) on ice for 30 min. The lysate was clarified at 16,000 RCF for 10 min at 4°C, and aliquoted into new tubes for storage at −80°C. Total protein concentration was determined by DC Protein assay using DC Protein Assay Reagents A, B, S (Bio-Rad, 5000113, 5000114, 5000115). The cell extracts were mixed with 4X LDS sample buffer (Life Technologies, B0007), 10X NuPAGE Sample Reducing Agent (Invitrogen, NP0009) and heated at 70°C for 10 min. The same amount of protein (20 μg) was loaded on NuPAGE gels (Life Technologies, NP0321). Protein lysates were separated on 4% to 12% Bis-Tris NuPAGE gels and run with MOPS Running Buffer (Life Technologies, J62847), transferred to polyvinylidene difluoride membrane (Millipore, IPVH00010), and blocked in 5% milk powder in TBST for 1 hr. The membranes were then incubated in primary antibody overnight with rocking at 4°C followed by incubation in appropriate secondary antibodies conjugated to horseradish peroxidase (1:5000) for 1 hr and detection by Super Signal West Pico PLUS Chemiluminescent Substrate (Thermo Fisher Scientific, 34577). Primary antibodies used were: Beclin-1 (Cell Signaling Technology, D40C5), LC3A/B (Cell Signaling Technology, D3U4C), Atg5 (Cell Signaling Technology, D5F5U), Atg12 (Cell Signaling Technology, D88H11), Atg16l1 (Cell Signaling Technology, D6D5), Atg7 (Cell Signaling Technology, D12B11), SQSTM1/p62 (Cell Signaling Technology, 5114), Cleaved Caspase-3 (Cell Signaling Technology, 5A1E), Ku80 (Invitrogen, PA5-17454), Rad51 (Cell Signaling Technology, D4B10), cMyc (Abcam, AB32072), Vinculin (Cell Signaling, 13901) and GAPDH (ProSci, 3783, 1:5000). 1:1000 dilution for primary antibodies was used unless otherwise indicated.

### Animal Experiments

Female athymic nude (Nu/Nu) mice with 6 weeks of age were purchased from Charles River Laboratory and used for subcutaneous engraftment. All the mice were housed in the Genome and Biomedical Sciences Facility at University of California, Davis under specific pathogen free environment, ambient temperature and a standard light-dark cycle. All the animal experiments were conducted in accordance with procedures approved by the Intuitional Animal Care and Use Committee (IACUC). For the subcutaneous engraftment, cells were trypsinized and resuspended in 50 µL of Matrigel (Fisher Scientific, CB-40230C). All the mice were either subcutaneously injected with 0.5 × 10^6^ cells of control or *Brca2*-KO KPC cells on the left and right flank, respectively. Following 3 days of tumor injection, the mice were randomized into 2 groups (8 mice per group). The mice were intraperitoneally administered with either vehicle or JQ1 (MedChemExpress, HY-13030) with a dose of 50 mg/kg 5 times per week. To prepare drug treatment for *in vivo* experiment, the vehicle consisted of 5% DMSO and 95% (20% SBE-β-CD in saline), while JQ1 was first resuspended in DMSO and then diluted with the vehicle to the final concentration of 5 mg/ml. Tumor size was measured with Vernier calipers three times a week, and tumor volumes were calculated according to the formula: 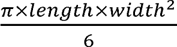 (15). After_6_ the course of drug administration, the mice were euthanized, and tumors were collected for histological examination. Note that one mouse in the DMSO group was lost, which is unrelated to the experiment.

Additional experimental details and methods can be found in the Supplementary information.

### Data Availability

The accession number for the data for RNA-seq and CUT-and-RUN-seq is NCBI GEO: GSE207298.

## Results

### Highthroughput drug screens in *Brca2*-deficient PDAC cells identify BET inhibitor

The rarity of FPC gene mutations in PDAC patients and commercially available PDAC cell lines hampered the development of personalized medicine for FPC patients (16). We reasoned that isogenic *Brca2*-deficient PDAC cells would enable us to identify unique vulnerabilities associated with *BRCA2* deficiency. To this end, we inactivated *Brca2* utilizing CRISPR/Cas9 in the murine PDAC cell lines derived from *Kras^+/LSL-G12D^; Trp53^+/LSL-R172H^; Pdx1-Cre* (KPC) mouse (herein after referred to as ‘KPC-mT3’ and ‘KPC-mT19’). (Fig. 1A and Supplementary Fig. 1A, B). As a control, we used a gRNA targeting *Rosa26* locus as described elsewhere (17). From this approach, we were able to establish the clonally derived isogenic *Brca2* knock-out (KO) KPC-mT3 and -mT19 cell lines. *Brca2*-KO KPC-mT3 cells exhibited a defect in RAD51 foci formation upon DNA damage and reduced HR efficiency in HR efficiency assay (Supplementary Fig. 1C-F). To determine if the isogenic *Brca2*-deficient KPC cell lines recapitulate the drug sensitivity observed in the clinical setting, we subjected our *Brca2*-KO KPC-mT3 cells to platinum-based chemotherapies and PARP inhibitors, which have been shown to be clinically effective in *BRCA1/2* mutant PDAC patients (8, 18). *Brca2*-KO KPC-mT3 cells displayed an increased sensitivity to oxaliplatin and PARPi, olaparib and talazoparib, compared to the control, without any proliferation change (Supplementary Fig. 1G-H). However, we did not see any significant change for other common first-line chemotherapies gemcitabine and 5-FU, suggesting that the increased sensitivity to PARPi is specifically due to *Brca2* deficiency. To identify new vulnerabilities in *BRCA2* deficient PDAC, we performed a HTS with a set of epigenetic drugs, clinically approved drugs, and 133 National Cancer Institute (NCI) approved drugs (Fig. 1A) as described previously (13). For screening a set of epigenetic drugs and 3300 approved drugs, we tested at a single concentration (2 µM) for both control and *Brca2*-KO KPC-mT3 cells (Supplementary file 1). 110 compounds that exhibited more than 50% reduced viability in either control or *Brca2*-KO KPC-mT3 cells were chosen for concentration response curve (CRC) experiments (Fig. 1B and Supplementary file 2). From the screening of a set of epigenetic drugs and 3300 approved drugs, we identified JQ1 as a top hit (Fig. 1B and Supplementary Fig. 2A). JQ1 rendered the biggest difference in IC50, indicating that *Brca2*-KO KPC-mT3 cells are preferentially sensitive to JQ1, a BET inhibitor. We further tested if JQ1 stood out in the second round of drug screening with the 133 NCI approved drug panel. This analysis confirmed that JQ1 is one of the top hits compared to the hits from the 133 NCI drug panel (Fig. 1C, Supplementary Fig. 2B-C and Supplementary file 3).

**Fig. 1.**
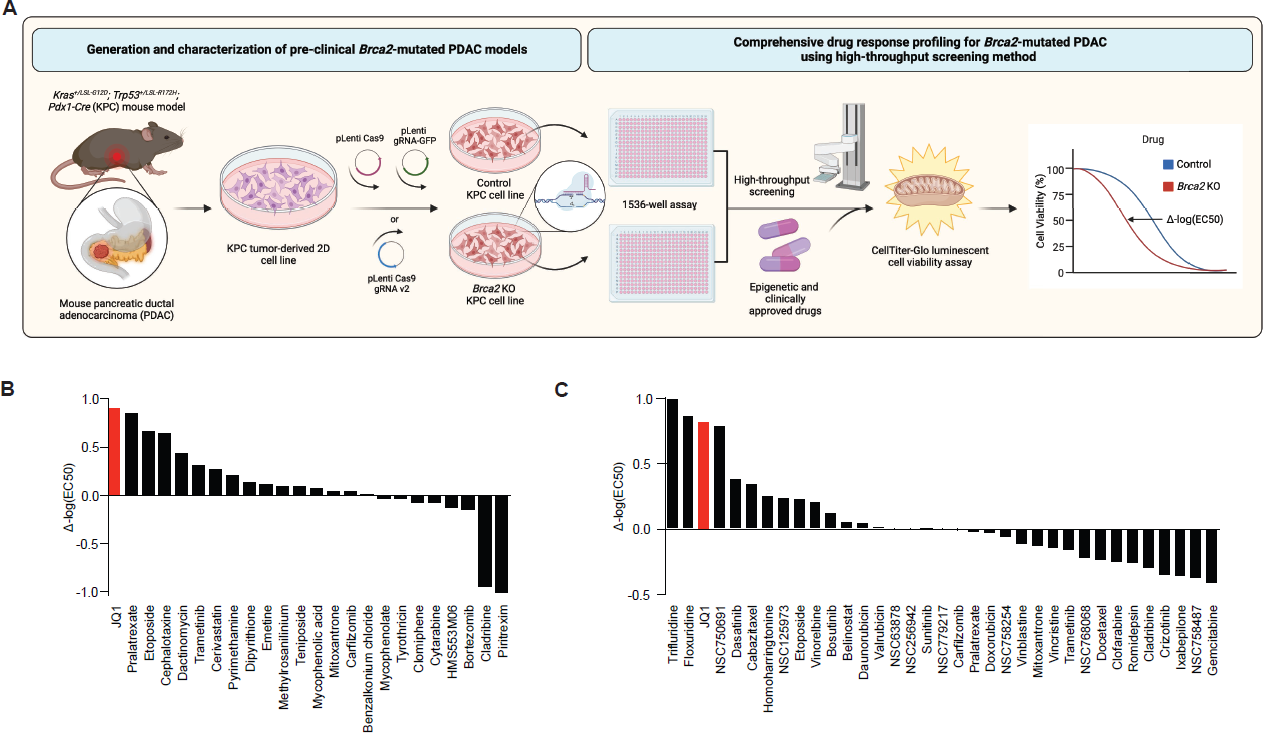
High-throughput screening (HTS) assays reveal sensitivity to BET inhibition in *Brca2*-deficient pancreatic cancer. **A,** Schematic illustration of the process to generate *Brca2* knock out (KO) murine *Kras^+/LSL-G12D^; Trp53^+/LSL-R172H^; Pdx1-Cre* (KPC) pancreatic cancer cell lines and HTS assays of a set of epigenetic durgs, clinically approved 3300 drugs, NCI approved 133 drugs with control and *Brca2*-KO KPC-mT3 cells. **B,** A set of epigenetic drugs and 3300 clinically approved drug screening results. **C,** FDA approved NCI 133 drug screening results.

### *Brca2*-deficient PDAC cells are sensitive to BET inhibition

To validate our HTS results, we tested JQ1 as well as birabresib and molibresib, additional BET inhibitors with two independent clonally derived *Brca2*-KO KPC cells from two different KPC cell lines each (KPC-mT3 and KPC-mT19-v2). We confirmed that *Brca2* deficiency rendered increased sensitivity to BET inhibitors (Fig. 2A-C). To confirm our findings in human PDAC cells, we additionally generated *BRCA2*-deficient MiaPaCa-2 human PDAC cell line and confirmed that *BRCA2* deficiency conferred increased sensitivity to BET inhibitors and olaparib (Fig. 2D and Supplementary Fig. 3A-B). This finding was further corroborated by the observation that CAPAN1, a *BRCA2* mutated PDAC cell line showed lower IC50 than three other cell lines with no mutation in *BRCA2* (Fig. 2E). Consistent with our finding in PDAC, a publicly available database for Genomics of Drug Sensitivity in Cancer (19) revealed that PARP inhibitors and BET inhibitors (e.g., JQ1, PFI-1, I-BET-762) are preferentially cytotoxic in mutant *BRCA2* context in breast cancer and pan-cancer, respectively (Fig. 2F**, G**). We additionally validated two other drugs, etoposide and afatinib from the HTS as these drugs have also been shown to be effective in *BRCA2* deficient setting (Supplementary Fig. 3C) (20, 21). Consistent with pharmacological BET inhibition, depletion of Brd2, Brd3 and Brd4 by transfection of pooled siRNA against Brd2, Brd3 and Brd4 in *Brca2*-KO cells resulted in a significant decrease in cell viability (Fig. 2H**, I** and Supplementary Fig. 3D, E). While BETi competitively blocks the bromodomain pocket of BET proteins, BETi has differential affinities to BRD2, BRD3, and BRD4 (22). For instance, JQ1 is known to bind with the highest affinity to BRD4 (23). To discern which BET protein is responsible for the increased sensitivity to BET inhibition in *Brca2*-KO KPC-mT3 cells, we individually knocked down Brd2, Brd3, and Brd4 using siRNA transfection. Brd4 depletion, but neither Brd2 nor Brd3, led to the reduced viability only in *Brca2*-KO KPC-mT3 cells, indicating that the inhibition of BRD4 function is critical for BET inhibition in *Brca2*-KO KPC-mT3 cells (Supplementary Fig. 3F, G). Previously, BET inhibition has been shown to down-regulate DDR genes and MYC expression, resulting in deleterious effects in PDAC (24, 25). While we observed similar down-regulation of DDR genes and MYC expression, these did not seem to be responsible for the differential response to BETi in *Brca2*-deficient KPC-mT3 cells (Supplementary Fig. 4A-B). In addition, BET inhibition was reported to induce BRCAness in HR-proficient PDAC cells, leading to the increased sensitivity to PARP inhibitor (24). The previously reported synergism between JQ1 and PARPi disappeared in *Brca2*-deficient KPC-mT3 cells, likely due to the increased sensitivity to BETi in *Brca2* deficiency (Supplementary Fig. 5).

**Fig. 2.**
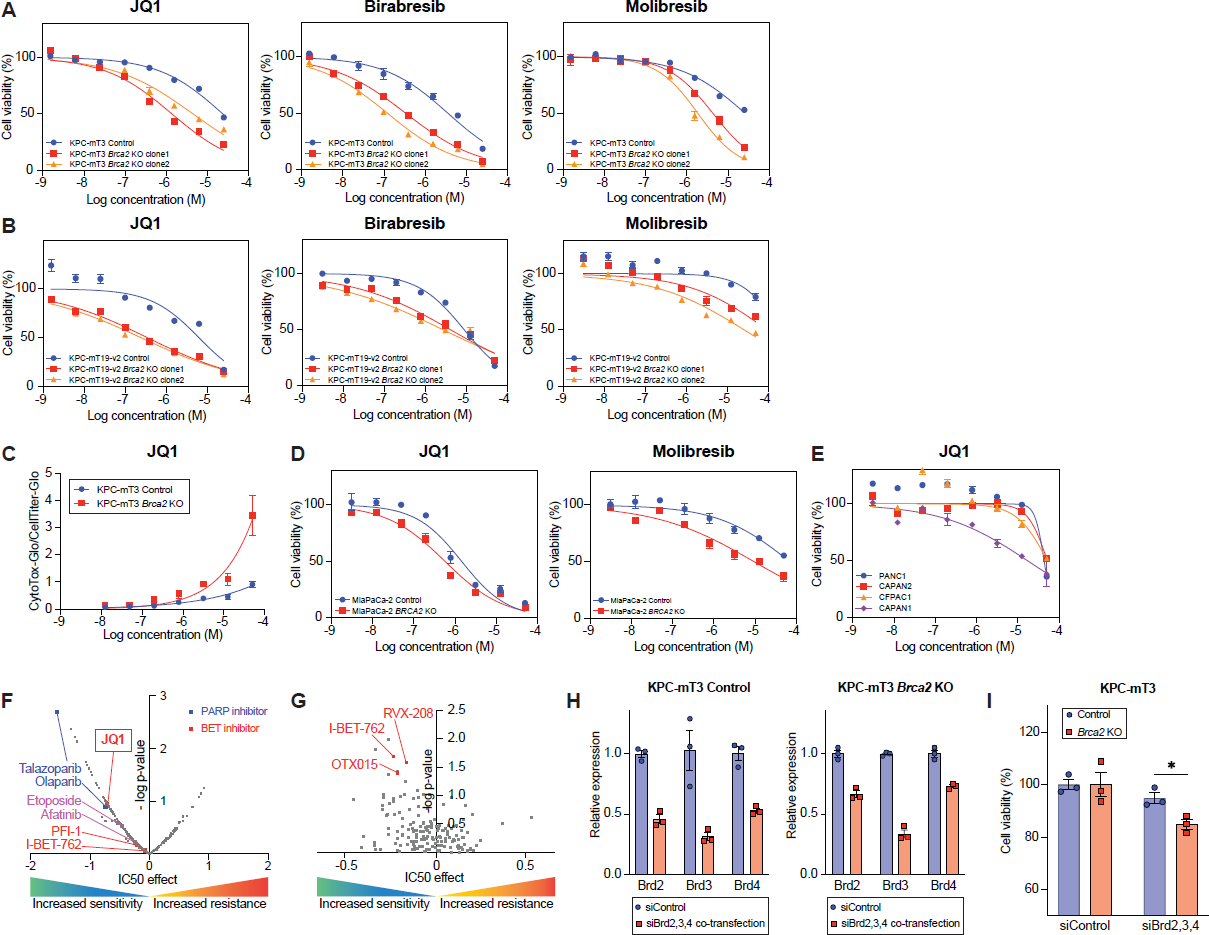
BRCA2 deficiency confers sensitivity to BET inhibitors. **A-B,** Cell viability assay with JQ1 and other BET inhibitors (birabresib and molibresib) in control and *Brca2*-KO KPC-mT3 (A) and -mT19 (B) cells. **C,** Cytotoxicity measured by CytoToxGlo assay in control and *Brca2*-KO KPC-mT3 cells upon JQ1 treatment. **D,** Cell viability assay with MiaPaCa2 control and *BRCA2*-KO cells upon BET inhibitiors treatment. **E,** Cell viability assay with the indicated cell lines upon JQ1 treatment. CAPAN1 is a *BRCA2* mutated PDAC cell line. **F**-**G**, Drug response data and *BRCA2* mutations in breast cancer (**F**) and pan-cancer (**G**) from the Genomics of Drug Sensitivity in Cancer. **H,** Knockdown efficiency upon siControl and pooled siBrd2, Brd3, and Brd4 transfection, determined by quantitative reverse transcription PCR (RT-qPCR) analysis. **I,** Percentage cell viability upon siControl or pooled siBrd2, Brd3, and Brd4 transfection. The mean ± SEM is shown. *p<0.05 was determined by one-tailed unpaired Student *t*-test.

### JQ1 significantly suppresses progression of *Brca2*-deficient KPC tumors *in vivo*

To evaluate if BET inhibition is an effective therapeutic approach for *BRCA2*-deficient pancreatic cancer *in vivo*, we subcutaneously injected KPC-mT3 cells into immunocompromised athymic nu/nu mice, and administered either with vehicle or JQ1 (50 mg/kg) via intraperitoneal injection (Fig. 3A). Over the 14-day treatment period, JQ1 treatment significantly reduced the tumor volume in both control and *Brca2*-deficient tumors, but more significant reduction in *Brca2*-deficient tumors (Fig. 3B-C). To evaluate the acute effect of JQ1 administration *in vivo*, we collected tumors after one-week treatment of JQ1 in separate cohorts of mice subcutaneously injected with control and *Brca2*-KO KPC-mT3 and mT19-v2 cells (one clone from KPC-mT3 and two clones from KPC-mT19-v2 with their control cells) (Fig. 3D). Immunohistochemical (IHC) analyses revealed that BET inhibition reduced cell proliferation and increased cell death, characterized by Ki67 IHC and TUNEL staining, respectively (Fig. 3E-J, Supplementary Fig. 6A-C). We did not see obvious evidence of cleaved caspase 3 activation (Supplementary Fig. 6D,E), suggesting alternative mechanisms of growth inhibition and cell death. Taken together, BET inhibition reduced the growth of PDAC tumors *in vivo* with more pronounced effect in *BRCA2* deficiency.

**Fig. 3.**
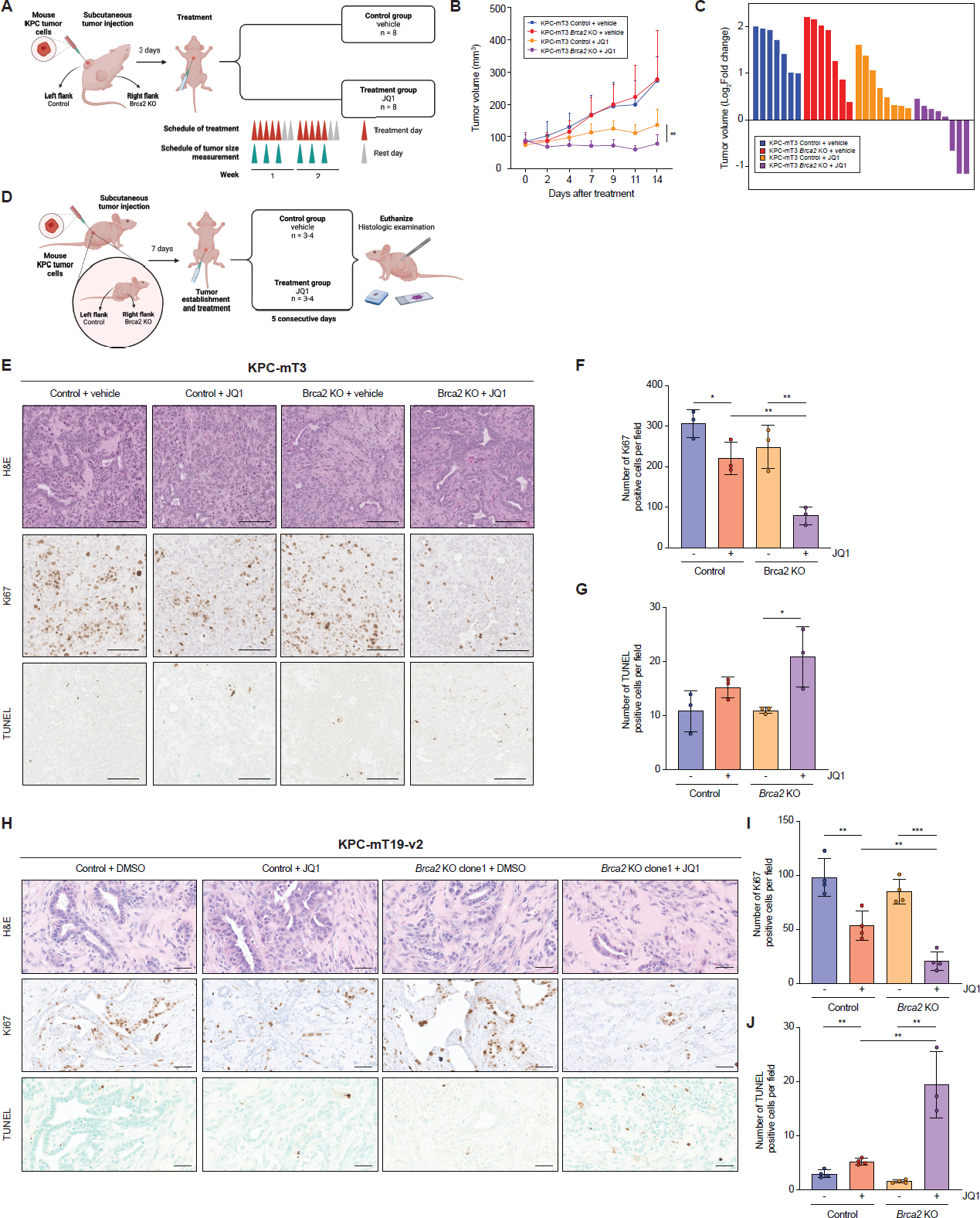
JQ1 treatment significantly suppresses *Brca2*-deficient pancreatic tumor growth *in vivo*. **A,** Schematic drawing of the experimental design. **B,** Mean tumor volume of the indicated groups (control+vehicle, n=7; control+JQ1, n=8; *Brca2*-KO+vehicle, n=7; *Brca2*-KO+JQ1, n=8). The mean±SD is shown. **p<0.05 was determined by two-tailed unpaired Student’s *t*-test. **C,** Changes in tumor volume after 14 days of JQ1 treatment. **D,** Schematic drawing of the experimental design to evaluate the acute response (7 days) of JQ1. **E-J,** Representative hematoxylin and eosin (H&E) staining (top), Ki67 IHC (middle) and TUNEL (bottom) staining of the indicated tumors (E and H) and quantifications of Ki67 (F and I) and TUNEL (G and J) positive cells. Scale bars, 100 µm. The mean±SD is shown. *p < 0.05, **p < 0.01, and ***p < 0.001 were determined by two-tailed unpaired Student’s *t*-test.

### *Brca2* deficiency augments JQ1-mediated upregulation of autophagy-related genes in PDAC

To extend our finding in *Brca2*-deficient PDAC cells, we tested BET inhibitors in *Atm*- and *Bub1b*-KO KPC-mT3 cells. Inactivation of these genes resulted in the increased sensitivity to BET inhibitors, suggesting that a defect in DNA damage response (DDR) or HR DNA repair pathway rendered a new vulnerability to BET inhibitors (Supplementary Fig. 7A-D). To elucidate the common mechanism behind the increased sensitivity to JQ1 in *Brca2*-KO, *Atm*-KO, and *Bub1b*-KO (hereinafter referred to as FPC-gene-KO) KPC-mT3 cells, we performed RNA-seq to compare the transcriptomic differences between FPC-gene-KO KPC-mT3 cells and the control upon JQ1 treatment. As expected, JQ1 drastically changed the transcriptomic profiles of the PDAC cells (Fig. 4A). Gene set enrichment analysis (GSEA) revealed that the genes involved in the epigenetic regulatory pathway were significantly upregulated upon JQ1 treatment (Fig. 4B). Interestingly, the gene sets related to macroautophagy and many autophagy-related pathways (e.g., phagosome acidification, phagosome maturation, and regulation of macroautophagy) were also significantly upregulated upon JQ1 treatment. The analysis of publicly available transcriptomic data (26, 27) from four different human PDAC cell lines with JQ1 treatment also confirmed the consistent up-regulation of the gene set involved in macroautophagy (Supplementary Fig. 8A, B). Macroautophagy is a proteo-homeostatic mechanism that sequesters and traffics unwanted proteins and/or organelles to the lysosome for degradation (28). Dysfunction in the autophagy process has been widely implicated in cancer, both as a cancer-promoting mechanism in times of nutrient deprivation and as an anti-cancer mechanism through autophagy-dependent cell death (28). When using GSEA to compare the transcriptomic changes between the control and FPC-gene-KO KPC-mT3 cells upon JQ1, we found that the up-regulation of macroautophagy and autophagy-related pathways were even more prominent in FPC-gene-KO KPC-mT3 cells (Fig. 4C-E), suggesting that the increased autophagic flux might play a role in the drug response of FPC-gene-KO KPC-mT3 cells. To determine if the changes to the level of autophagy-related transcripts upon JQ1 were regulated at the epigenetic level, we performed CUT-and-RUN-seq for H3K27ac to identify the genomic regions of the active promoters and enhancers in the control and *Brca2*-KO KPC-mT3 cells upon JQ1 treatment. The H3K27ac CUT-and-RUN-seq analysis showed that the autophagy-related genes that were up-regulated upon JQ1 treatment had increased H3K27ac in the proximal transcriptional start site in the JQ1 treated *Brca2*-KO KPC-mT3 cells compared to the control (Fig. 4F,G), suggesting that *Brca2*-KO KPC-mT3 cells might exhibit more autophagy and JQ1 might preferentially induce autophagy mediated cell death in *Brca2*-KO KPC-mT3 cells. Taken together, JQ1 transcriptionally upregulates the genes associated with autophagy in *Brca2*-deficient PDAC.

**Fig. 4.**
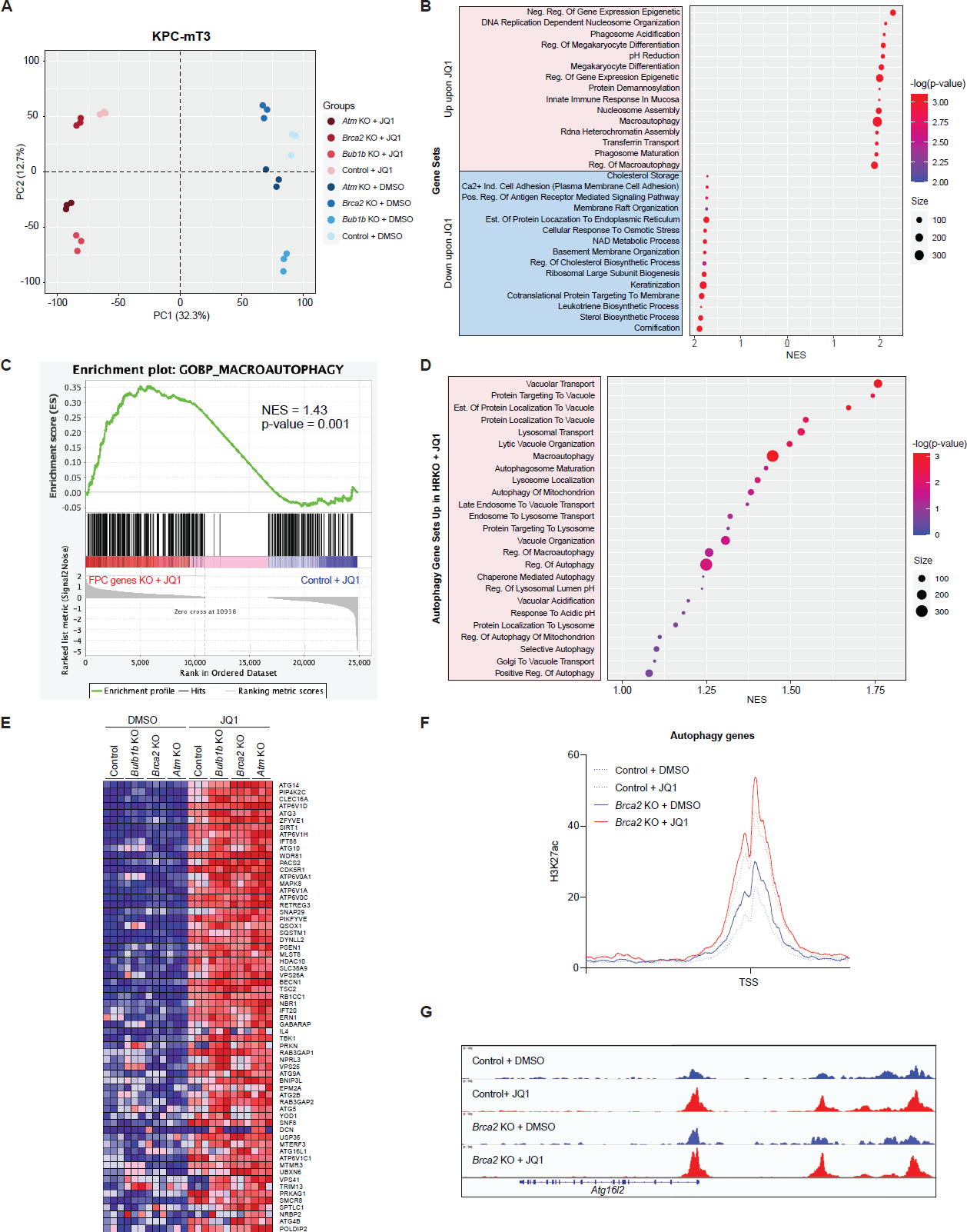
JQ1 significantly upregulates autophagy-related processes in FPC-gene-KO KPC cells. **A,** Principal Component Analysis of the transcriptome of FPC-gene-KO KPC cells (*Atm*-KO, *Brca2*-KO, *Bub1b*-KO) and the control, treated with 1 µM JQ1 or DMSO for 72 hrs (n=3). **B,** Top 15 upregulated and downregulated gene sets from Gene Set Enrichment Analysis (GSEA) with Gene Ontology (GO) Biological Processes (BP) gene sets upon treatment with JQ1 (n=3). The size and the color of each point indicate the number of genes and the significance of each output as −log(p-value), respectively. **C,** GSEA of GO BP Macroautophagy gene set comparing JQ1-treated FPC-gene-KO KPC cells and JQ1-treated control KPC cells. **D,** Top autophagy-related gene sets from GSEA of transcriptomic profiles of JQ1 treated FPC-gene-KO cells, and JQ1 treated control (n=3). **E,** A heatmap of differentially expressed autophagy-related genes. **F-G,** A metaprofile of CUT-and-RUN H3K27ac marks proximal to transcriptionally upregulated autophagy genes in the macroautophagy gene set (F) and a representative browser track image in *Atg16l2* locus (G). NES, Normalized Enrichment Score. TSS, Transcription Start Site.

### *Brca2* deficiency increases autophagic flux upon JQ1 treatment

From the transcriptomic analyses, we hypothesized that JQ1-mediated upregulation of autophagy-associated genes induces increased autophagic flux in *Brca2*-KO KPC cells. Recent reports suggested that JQ1 induces the accumulation of autophagosomes and autolysosomes, and increased autophagy flux (29). To test this hypothesis, we measured the expression of autophagy protein markers upon JQ1 treatment. *Brca2*-KO KPC-mT3 cells exhibited higher base-line expression of autophagy markers LC3A/B II and BECLIN-1 compared to the control, and JQ1 treatment increased the expression LC3A/B II and BECLIN-1 in both control and *Brca2*-KO KPC-mT3 cells (Fig. 5A). Particularly, JQ1-treated *Brca2*-KO KPC-mT3 cells had further increased expression of autophagy marker proteins along with decreased expression of autophagy substrate SQSTM1/p62, indicative of increased autophagic flux. Despite an increase in autophagic flux in JQ1-treated *Brca2*-KO KPC-mT3 cells, there was no significant difference in the expression of ATG family proteins (Supplementary Fig. 9). Immunofluorescence (IF) staining of LC3 showed more autophagosome formations depicted by the increased number of LC3 puncta in *Brca2*-KO cells upon JQ1 treatment (Fig. 5B**, C**). To visualize the autophagy flux at the cellular level, we utilized an autophagy reporter construct (EGFP-LC3-RFP) to observe autophagosome puncta formation and trafficking in the cell. This reporter allowed us to distinguish early autophagic vacuoles from autolysosomes using the differential pH sensitivities as described elsewhere (30). JQ1 treatment in control and *Brca2*-KO KPC-mT3-v2 cells increased LC3 puncta intensity per cell (Fig. 5D**, E**). Consistent with IF results, *Brca2*-KO KPC-mT3-v2 cells also had higher baseline LC3 puncta intensity compared to control cells, and this intensity in the *Brca2*-KO KPC-mT3-v2 cells further increased upon JQ1 treatment. Using the recently developed quantitative method to measure dynamic autophagy rates in live cells (31), we measured the rates of three major steps in autophagy: the rate of formation of autophagosomes (R_1_), the rate of fusion of autophagosomes with autolysosomes (R_2_), and the rate clearance of autolysosomes (R_3_) (Fig. 5F). First, we validated our system with rapamycin, an autophagy inducer, resulting in the increased R_1_, R_2_, and R_3_ of murine KPC control cells (Supplementary Fig. 10A). The measurement of dynamic autophagy rates revealed that *Brca2*-KO KPC-mT3-v2 cells have higher basal autophagy flux (Fig. 5G), and JQ1 treatment further increased autophagy flux in both control and *Brca2*-KO cells (Fig. 5H, Supplementary Fig. 10B). Collectively, we concluded that JQ1 induces increased autophagic flux in *Brca2*-deficient PDAC cells.

**Fig. 5.**
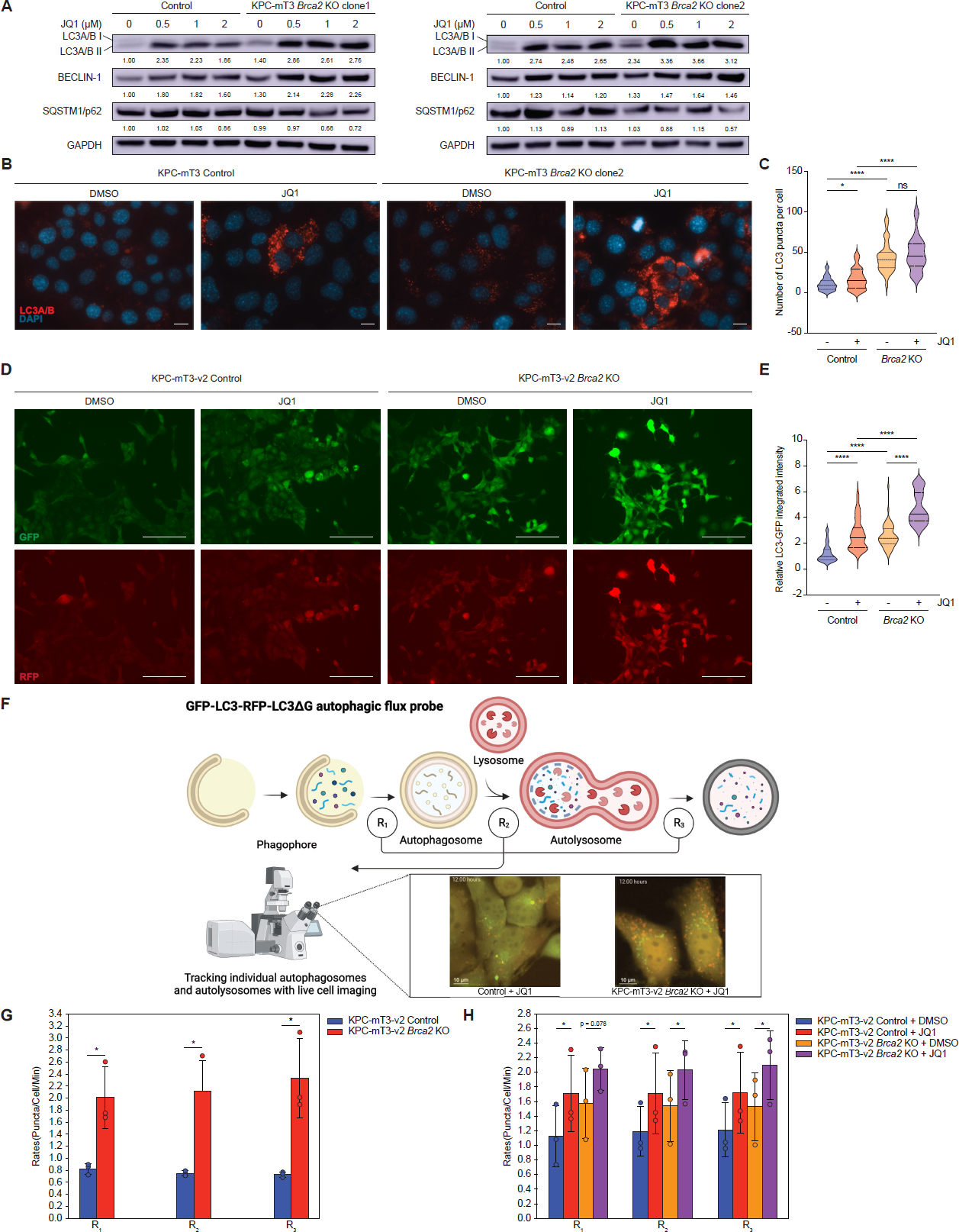
JQ1 and *Brca2* deficiency increase autophagic flux. **A,** Western blotting of autophagy markers (LC3A/B, BECLIN-1, and SQSTM1/p62) in control and *Brca2*-KO KPC-mT3 clone 1 and 2 upon JQ1 treatment for 72 hrs. **B**-**C,** Representative images of immunofluorescence staining of LC3 upon 1 μM JQ1 treatment for 24 hrs (**B**) and the quantification (**C**) in control and *Brca2*-KO KPC-mT3 cells. Scale bars, 100 µm. **D** and **E**, Autophagic flux with the autophagy reporter (EGFP-LC3-RFP) upon 5 μM JQ1 treatment for 24 hrs in control and *Brca2*-KO KPC-mT3-v2 cells. *p<0.01, ****p<0.0001, and ns, not significant were determined by two-tailed unpaired Student’s *t*-test. Scale bars, 150 µm. **F,** A schematic drawing of the live cell imaging-based autophagy rate measurement. **G,** Basal rates of *Brca2*-KO and control KPC-mT3-v2 cells. A minimum of 600 cells were imaged. Bar graphs represent mean with error bars representing ± standard deviation based on three biological replicates. *p<0.05, and ns, not significant were determined by one tailed paired *t*-test. **H,** Rates of *Brca2*-KO and control KPC-mT3-v2 cells after treatment with 5 µM JQ1 for 12 hours. A minimum of 600 cells were imaged. Bar graphs represent mean with error bars representing ± standard deviation based on three biological replicates. *p<0.05, and ns, not significant were determined by one-tailed paired *t*-test.

### BET inhibition in *Brca2* deficiency causes autophagy-dependent cell death

The increased autophagy induced by JQ1 in *Brca2*-KO KPC-mT3 cells prompted us to investigate whether autophagy-dependent cell death is critical for *Brca2*-KO KPC-mT3 cells upon JQ1 treatment. Previously, JQ1 has also been known to induce ferroptosis, an iron-dependent cell death (32). However, the increased sensitivity to JQ1 in *Brca2*-KO KPC-mT3 cells was not due to increased ferroptosis (Supplementary Fig. 11A, B). Furthermore, cleaved caspase 3 expression was not significantly increased in *Brca2*-KO KPC-mT3 cells upon JQ1 treatment, compared to the control cells treated with JQ1 (Supplementary Fig. 11C). This indicated that the preferential cell death in *Brca2* deficiency is not likely due to either ferroptosis or caspase3-dependent apoptosis. To show JQ1-mediated cell death in *Brca2*-KO KPC-mT3 cells is autophagy-dependent, we asked whether hydroxychloroquine (HCQ), an autophagy inhibitor, can rescue JQ1-induced cell death in *Brca2*-KO KPC-mT3 cells. As expected, HCQ treatment partially rescued the increased sensitivity to JQ1 in *Brca2*-KO-mT3 cells while the control cells did not show any antagonistic effect upon JQ1 and HCQ treatment (Fig. 6A**, B** and Supplementary Fig. 11D). Likewise, we were able to rescue the increased sensitivity to JQ1 with autophagy inhibition in *Atm*- and *Bub1b*-deficient KPC cells, suggesting that autophagy-dependent cell death might be a common mechanism in HR-deficiency (Fig. 6C**, D**). While HCQ is known to block the fusion of autophagosomes with lysosomes, it also has other autophagy-independent effects. Therefore, we decided to genetically perturb the autophagy genes responsible for autophagosome formation such as *Atg5, Atg12* and *Atg16l1* using RNA interference. In line with pharmacological inhibition of autophagy, depletion of autophagy genes also rescued the effect of JQ1 in *Brca2*-KO KPC-mT3 cells (Fig. 6E**, F** and Supplementary Fig. 11E). Taken together, our data with pharmacological and genetic perturbation of autophagy indicated that the induction of autophagy-dependent cell death in *Brca2* deficiency is in part responsible for the increased sensitivity to BET inhibition in *Brca2*-KO setting.

**Fig. 6.**
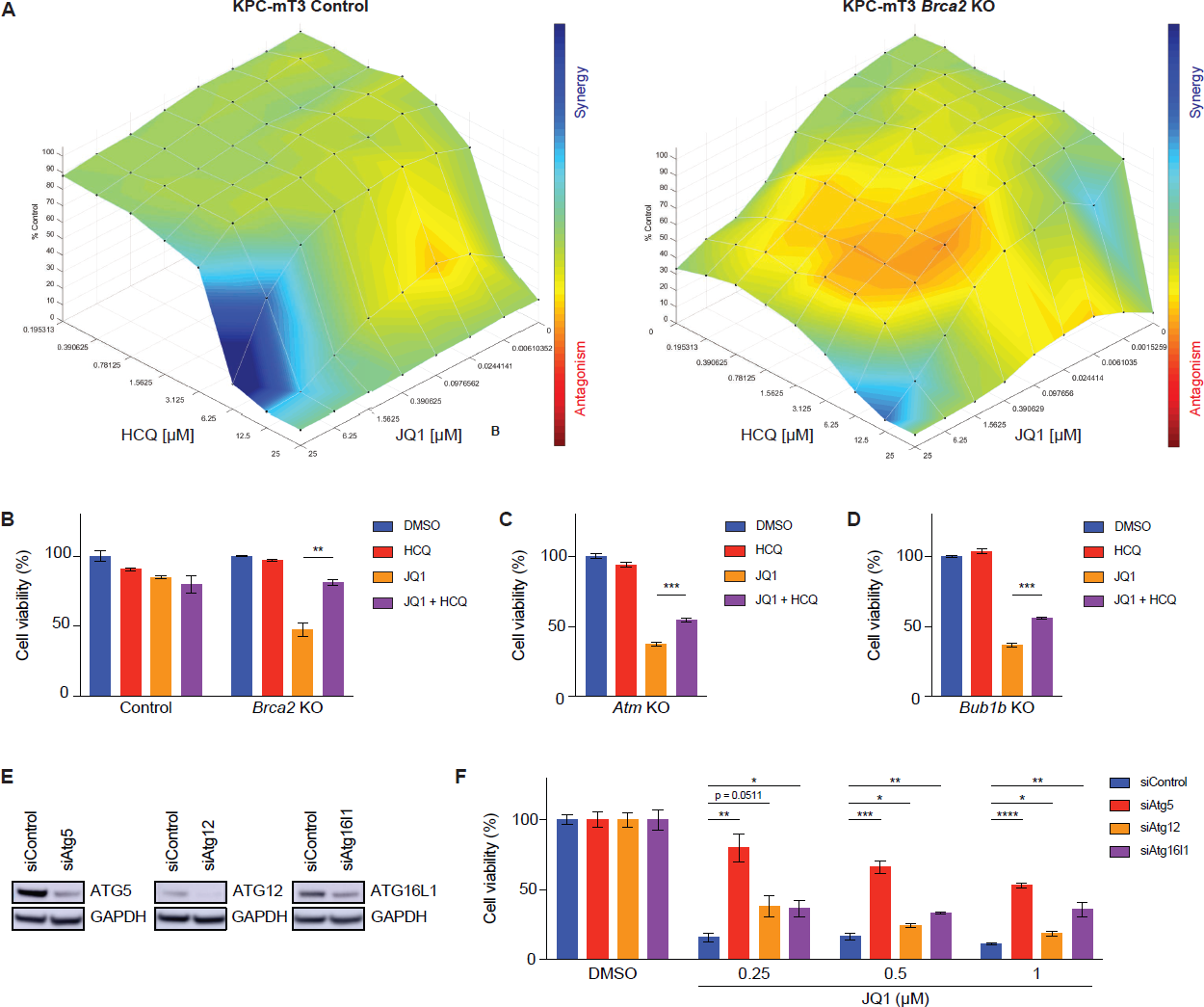
Autophagy inhibition partially rescues JQ1-mediated cell death in *Brca2*-KO KPC cells. **A,** Analysis and visualization of hydroxychloroquine (HCQ) and JQ1 combination of control (left) and *Brca2*-KO KPC-mT3 cells (right, clone 2) with Combenefit. Blue and red indicate synergy and antagonism, respectively. **B, C,** and **D,** Cell viability assay of control, *Brca2*-KO (**B**), *Atm*-KO (**C**) and *Bub1b*-KO (**D**) KPC-mT3 cells upon DMSO, HCQ (5 μM), JQ1 (0.5 μM), HCQ (5 μM) and JQ1 (0.5 μM) combination treatment. **E,** Western blotting of *Brca2*-KO KPC-mT3 cells transfected with the indicated siRNAs. **F,** Percentage cell viability of *Brca2*-KO KPC-mT3 cells transfected with the indicated siRNAs upon JQ1 treatment. Cell viability was calculated as the average of the percentage covered area per field (see Supplementary Fig. 11E). *p<0.05, **p<0.01, ***p<0.001 and ****p<0.0001 were determined by two-tailed unpaired Student’s *t*-test.

## Discussion

In this study, we identified JQ1, a small molecule BET inhibitor from the high-throughput drug screen in *Brca2*-KO KPC cells. This finding was validated using isogenic human and mouse cell lines, human PDAC cell lines and publicly available pan-cancer analysis. We propose that BET inhibition might be a potential therapeutic strategy for FPC patients with the mutated genes involved/implicated in HR DNA repair pathway. BET proteins have recently emerged as therapeutic targets for human malignancies, and BRD4 among BET proteins appears to be a major target of BET inhibition (33). In line with this, our genetic perturbation experiments using siRNA confirmed that the increased sensitivity to BET inhibitors in *Brca2*-KO KPC cells was mainly mediated by BRD4. While the BET proteins share ∼75% identity between family members, it has been shown that they play distinct roles in gene transcription, chromatin remodeling, DNA replication, and DNA damage repair (9). In particular, a recent study further confirms that BRD4 is directly involved in double strand DNA repair (34). In addition, due to a poor pharmacokinetic profile, low oral bioavailability, and unwanted toxicities of JQ1, there has been extensive effort to develop selective inhibitors of BET with reduced toxicity. Nonetheless, our findings highlight a novel vulnerability of FPC, particularly in *BRCA2* deficiency. Identification of BET inhibitors as a new vulnerability of FPC was an unexpected finding because many epigenetic drugs including BET inhibitors were used to induce a “BRCAness” phenotype and exert a synergistic cytotoxicity effect in combination with olaparib in HR-proficient tumors, including breast, ovarian and prostate cancers (35). This phenomenon has also been observed in PDAC upon combination therapy with JQ1 and olaparib as JQ1 mitigates double stranded break repair pathway and further improves the efficacy of olaparib (24). However, in our study, we found that *BRCA2*-deficient PDAC is sensitive to BET inhibition alone, which sheds light on unexplored mechanisms between HR DNA repair pathway and epigenetic regulations. Several PARP inhibitors have already been approved for *BRCA1/2* mutated cancers. However, the development of PARPi resistance appears to be inevitable (36). JQ1 has been shown to re-sensitize *BRCA2*-mutated ovarian cancer cells that have developed resistance to olaparib (37). It remains to be determined whether BETi can be still effective to *BRCA2*-mutated PDAC cells that become resistant to PARPi. Therefore, BET inhibition as a therapeutic approach might be broadly applicable to both HR-proficient and -deficient as well as PARPi-resistant cancers.

There is ample evidence that autophagy and DNA repair pathways are interconnected. For instance, autophagy deficiency has been shown to induce genomic instability through various mechanisms, reviewed in Vesonni et al. (38). Here, we showed that the increased sensitivity to BETi in *Brca2*-deficient PDAC cells is in part due to autophagy-dependent cell death. Autophagy is a conserved catabolic process that contributes to cellular homeostasis, through the degradation and recycling of cytoplasmic components and organelles in the lysosome (28, 39, 40). Recently, it has been reported that PDAC tumors exhibit high autophagy activity which confers resistance to chemotherapies (41). Therefore, a significant effort to combine autophagy inhibitor with other therapies are being pursued in both preclinical and clinical settings. For example, the combination of autophagy inhibitors with MAPK pathway inhibitors are currently being tested in patients with PDAC (41, 42, 43). We found that *Brca2* deficiency resulted in increased autophagy activity. This is in line with recent observations that *BRCA1/2* can negatively regulate autophagy (44, 45, 46). Upon BET inhibition, the autophagy activity was further pronounced, resulting in autophagy-dependent cell death. Unlike the current therapeutic strategy of combining autophagy inhibition with other chemo- or targeted therapies, autophagy inhibition confers antagonistic effect of JQ1 sensitivity in *Brca2* deficiency. This raises a potential concern that autophagy inhibition might exert antagonistic effect depending on the context. It should be noted that autophagy has been shown to have dual roles in cancer progression: both pro- and anti-tumorigenic roles (47, 48). For instance, suppression or deficiency of autophagy genes has been shown to enhance tumorigenesis. In pancreatic cancer, *Atg5* or *Atg7* deficiency resulted in increased PanIN lesions, albeit lack of progression to malignant disease (49). In line with this, a monoallelic deletion of *BECLIN1* is frequently found in breast cancer samples (50).

It is tempting to speculate that the increased sensitivity to JQ1 might be associated with DDR defects as we showed the defects in other FPC genes (e.g., *ATM, BUB1B*) resulted in increased sensitivity to BETi. Because of the scarcity of non-*BRCA* mutations in FPC, the clinical benefit of PARPi and other therapeutic strategies have only been focused on pancreatic cancer with *BRCA1/2* mutations. It is urgently required to develop appropriate preclinical models to evaluate the PARPi therapies and identify novel vulnerabilities tailored toward each FPC gene. Isogenic FPC gene KO of murine KPC or human PDAC cell lines can provide a platform to identify a new vulnerability of FPC and investigate the underlying molecular mechanisms of synthetic lethality and potential resistance mechanisms in the future.

## Acknowledgements

This study was supported by a grant from the National Cancer Institute of the National Institutes of Health 5K22CA226037 (to C.-I.H) and 5R37CA249007. C.-I.H was supported by the Synthego Genome Engineer Innovation Award. K.Y.J was supported by UC Davis Maximizing Access to Research Careers (MARC) Program (NIGMS-MARC-U-STAR grant T34GM136469), UC Davis College of Biological Sciences Dean’s Circle Summer Undergraduate Research Program, UC Davis Provost’s Undergraduate Fellowship, and Barry M. Goldwater Scholarship. S.A. was supported by the Anandamahidol Foundation (Thailand). We would like to thank Vincenzo Corbo, Rafaella Casolino, Sarah Wang, and Jihao Xu for their critical comments on the manuscript. We would like to thank the Hunter lab for providing immunofluorescence microscopy imaging, Sarah Wang for developing an ImageJ plugin that was used for quantifying LC3 puncta measurement. We gratefully acknowledge the support for histology and immunohistochemistry from the Center for Genomic Pathology at University of California Davis. Figures were generated with BioRender.com.

