## Supplementary information for "A new vulnerability to BET inhibition due to enhanced autophagy in BRCA2 deficient pancreatic cancer"

**Supplementary File 1. Data associated with 3300 clinically approved drug screening in *Brca2*-KO KPC and control cells**

**Supplementary File 2. Concentration Response Curve (CRC) data associated with 110 clinically approved drugs**

**Supplementary File 3. Concentration Response Curve (CRC) data from the 133 NCI drug panel**

**Supplementary File 4. Sequence information of the guide RNAs used in this study**

**Supplementary Figure Legends**

**Supplementary Fig. 1 Characterization of *Brca2*-KO KPC cells. A,** Surveyor assay performed with *Brca2* knock-out (KO) KPC-mT3 and mT19 cells to verify CRISPR/Cas9-mediated mutation. – and + indicates the absence and presence of T7 endonuclease I. **B,** Graph illustrating the Inference of CRISPR Edits (ICE) KO score for each *Brca2* KO KPC cell lines generated. **C-D,** Immunofluorescence (IF) staining of RAD51 foci in control and *Brca2*-KO KPC-mT3 cells upon 0.2 μM doxorubicin treatment for 24 hrs (C) and the quantifications of the number of RAD51 foci / cell (D). Scale bars, 100 µm. **E-F**, Homologous recombination efficiency assay in control and *Brca2*-KO KPC-mT3 cells (E) and the quantification (F). **G,** Cell viability assay performed with *Brca2*-KO KPC-mT3 cells upon 72 hrs treatment with various concentrations of PARP inhibitors (olaparib and talazoparib) and common first-line PDAC chemotherapies (gemcitabine, 5-FU and oxaliplatin). **H,** Relative cell growth of KPC-mT3 cells with control and *Brca2*-KO. Error bars represent mean ± SEM.

**Supplementary Fig. 2 *Brca2*-KO KPC cells are sensitive to JQ1.** **A,** Scatter plot analysis of the set of epigenetic drugs and clinically approved 3300 drugs screen. All data from all assay plates, including high and low controls, are displayed. The average signal-to-background ratio (S:B) = 123.67 ± 3.97 for control cells (left) and 109.36 ± 4.05 for *Brca2*-KO KPC-mT3 cells (right) (n = 3 plates). The heat cutoff was determined by using the Average + 3 x Standard Deviation of the low control. Z’ = 0.74 ± 0.04 for control cells and 0.53 ± 0.11 for *Brca2*-KO KPC-mT3 cells between high and low controls wells, as well as broad distribution of hits indicate that the assay is robust. Blue circles represent low control wells containing cells in the presence of DMSO only, black circles represent data wells containing compounds and red circles represent high control wells containing media + DMSO. **B,** Scatter plot analysis of NCI approved 133 drugs screen. All data from all assay plates, including high and low controls, are displayed. The average signal-to-background ratio (S:B) = 137.39 ± 5.61 for control cells (left) and 130.76 ± 5.61 for *Brca2*-KO KPC-mT3 cells (right) (n = 3 plates). The heat cutoff was determined by using the Average + 3 x Standard Deviation of the low control. Z’ = 0.75 ± 0.03 for control cells and 0.66 ± 0.08 for *Brca2*-KO KPC-mT3 cells between high and low controls wells, as well as broad distribution of hits indicate that the assay is robust. Blue circles represent low control wells containing cells in the presence of DMSO only, black circles represent data wells containing compounds and red circles represent high control wells containing media + DMSO. **C,** JQ1 concentration response curve of control and *Brca2*-KO KPC-mT3 cells in 1536 wells. Each curve represents the mean and the standard deviation of 16 replicates.

**Supplementary Fig. 3 New vulnerabilities of *BRCA2*-deficient pancreatic cancer. A,** Surveyor assay performed with control and *BRCA2*-KO MiaPaCa-2 cells to verify CRISPR/Cas9 mediated mutation. – and + indicates the absence and presence of T7 endonuclease I. **B,** Cell viability assay performed with control and *BRCA2*-KO MiaPaCa-2 cells upon olaparib treatment. **C,** Cell viability assay performed with control and *Brca2*-KO KPC-mT3 cells upon 72 hrs treatment with various concentration of afatinib (left) and etoposide (right). **D**-**E,** Cell viability of control and *Brca2*-KO KPC-mT3 cells were assessed after pooled transfection of siBrd2, siBrd3 and siBrd4 with the final concentration of 60 nM (**D**) and 75 nM (**E),** respectively. **F,** Knockdown efficiency upon siControl transfection or individual Brd2, Brd3, and Brd4 siRNAs, assessed by RT-qPCR. **G,** Percentage cell viability upon siControl transfection or individual Brd2, Brd3, and Brd4 siRNAs. The mean ± SEM is shown. *p<0.05, **p<0.01 and ***p<0.001 were determined by one-tailed unpaired Student *t*-test.

**Supplementary Fig. 4 JQ1 treatment down-regulates MYC and genes involved in DNA repair**. **A,** Western blot analysis of MYC, KU80 and RAD51 in *Brca2*-KO KPC-mT3 clone 1 (left) and 2 (right) and control from KPC-mT3 cells upon 0, 0.5, 1, and 2 μM of JQ1 treatment for 72 hrs. **B,** Western blot analysis of MYC in *Brca2-*KO KPC-mT3 clone 1 (left) and 2 (right) and control after 4 hrs and 24 hrs treatment of JQ1. GAPDH is the loading control.

**Supplementary Fig. 5 *Brca2* deficiency abrogates JQ1’s synergistic interaction with olaparib**. **A-B,** Analysis and visualization of olaparib and JQ1 in control (**A**) and *Brca2*-KO KPC-mT3 cells (**B**) with Combenefit. Blue indicates synergy and red indicates antagonism.

**Supplementary Fig. 6 JQ1 treatment is preferentially effective in *Brca2*-deficient tumors. A,** Representative hematoxylin and eosin (H&E) staining, Ki67 immunohistochemistry, and TUNEL staining of the indicated tumors derived from control and *Brca2*-KO clone 2 KPC-mT19-v2 cells. **B-C,** Quantifications of the number of Ki67 (B) and TUNEL (C) positive cells. **D-E**, Immunohistochemical staining of cleaved caspase 3 (D) and the quantification (E) in the indicated tumors derived from control and *Brca2*-KO KPC-mT3 cells. Error bars indicate mean ± SD. Statistical significance was analyzed by two-tailed unpaired Student’s *t*-test for histological analyses. *p < 0.05, **p < 0.01, ***p < 0.001, and ns, not significant. Scale bars, 100 µm.

**Supplementary** **Fig. 7 Characterization of *Atm*- and *Bub1*-KO KPC-mT3 cells. A,** Surveyor assay performed with *Atm and Bub1b* knock-out (KO) KPC-mT3 cells to verify CRISPR/Cas9-mediated mutation. – and + indicates the absence and presence of T7 endonuclease I. **B,** Graph illustrating the Inference of CRISPR Edits (ICE) KO score. **C-D,** Cell viability assay performed with *Atm*-KO (C) and *Bub1b*-KO (D) KPC-mT3 cells upon JQ1, birabresib and molibresib treatments.

**Supplementary Fig. 8 JQ1 treatment in human PDAC cells up-regulates the genes involved in autophagy. A,** Genes involved in macroautophagy were up-regulated upon JQ1 treatment in GSEA with two human PDAC cell lines, ASPC1 and PANC1 (GSE124069). **B,** Genes involved in macroautophagy were up-regulated upon JQ1 treatment in GSEA with two human PDAC cell lines, MiaPaCa-2 and L37PL in GSEA (GSE98067).

**Supplementary Fig. 9 JQ1 treatment increases autophagy in *Brca2*-KO KPC cells.** Western blot analysis of autophagy-related proteins (ATG5, ATG7, ATG12, ATG16L1) in *Brca2*-KO KPC-mT3 clone 1 and 2 and control upon 0, 0.5, 1, and 2 μM of JQ1 treatment for 72 hrs. GAPDH is the loading control from **Fig. 5A.**

**Supplementary Fig. 10 Quantitative measurement of dynamic autophagy rates. A,** Rates of KPC-mT3-v2 control cells after treatment with 100 nM Rapamycin for 30 minutes. Respective concentration of DMSO was used a negative control. A minimum of 600 cells were imaged. Bar graphs represent mean with error bars representing ± standard deviation based on three biological replicates. **B,** Autophagosome (AP) and autolysosome (AL) numbers were measured over 24 hrs following treatment with JQ1 or DMSO in control and *Brca2*-KO KPC-mT3-v2 cells. Values were normalized to AP and AL numbers at 0 hrs post-treatment treatment (AP_0_ and AL_0_). Line represents mean with shaded area representing ± standard deviation based on two biological replicates. Statistical significance was analyzed by one-tailed paired Student’s *t*-test. *p < 0.05, ns, not significant.

**Supplementary Fig. 11 Autophagy inhibition partially rescues JQ1-mediated cell death in *Brca2*-KO KPC cells.** **A,** Cell viability assay of *Brca2*-KO KPC-mT3 cells upon DMSO, ferrostatin (ferroptosis inhibitor, 5 μM), JQ1 (0.5 μM), JQ1 (0.5 μM) and ferrostatin (5 μM) combination treatment for 72 hrs. **B,** Cell viability assay of *Brca2*-KO KPC-mT3 cells upon DMSO, ferrostatin (5 μM), erastin (ferroptosis activator, 5 μM), ferrostatin (5 μM) and erastin (5 μM) combination treatment for 72 hrs. **C,** Western blot of cleaved caspase 3 in control and *Brca2*-KO KPC-mT3 cells upon DMSO, doxorubicin (0.2 μM), and JQ1 (10 μM) treatment for 48 hrs. VINCULIN is the loading control. **D**, Analysis and visualization of HCQ and JQ1 combination of *Brca2*-KO KPC-mT3 cells (clone 1) with Combenefit. Blue indicates synergy and red indicates antagonism. **E,** Representative images of control siRNA or Atg5 siRNA, Atg12 siRNA, Atg16l1 siRNA expressing *Brca2*-KO KPC-mT3 cells treated with JQ1. Microscopic images were taken from 4X field to quantify the covered area per field. *p<0.05, **p<0.01, ***p<0.001 were determined by two-tailed unpaired Student’s *t*-test. Scale bars, 800 µm.

**Supplementary Methods**

**Compound Library and HTS**

A collection of ~3,300 clinically approved drugs obtained from multiple vendors were assembled at the UF Scripps Biomedical Research High-throughput Screening Center and reformatted into 1536-well source plates for automated robotics screening (1). In addition, the NCI-approved oncology drug set of 133 compounds was obtained directly from the NCI, described previously (2, 3). In brief, a 1536-well 2D cell viability assay was optimized previously (2). Cell viability was determined based on the amount of ATP present using CellTiter-Glo (Promega, G7573). Trypsinized cells were resuspended and filtered through cell strainer. 62 cells in 5 µL of culture media were seeded in 1536-well plates. After incubation of the assay plates overnight, cells were treated with compounds and vehicle (10 nL, 0.02% DMSO). Cell viability was assessed after 72 hrs of incubation using CellTiterGlo reagent according to manufacturer’s instructions. All data files obtained were uploaded into Scripps’ institutional database for individual plate quality control and hit selection. All plates were required to pass Z’ analysis >0.5 based on the control wells (n =24 per control per plate) before they were subjected to further analysis (4). Compound activity was normalized on a per-plate basis as described previously. The High control is defined as wells containing medium only (100% inhibition), and Low control wells contain cells treated with DMSO only (0% inhibition). IC50 values were determined by fitting the concentration-response curve (CRC) data with a four-parameter variable-slope method in GraphPad Prism (GraphPad Software).

**Lentiviral and Retroviral Transduction**

To produce the lentivirus and retrovirus, HEK-293T (for lentivirus production) or Phoenix Ecotropic (for retrovirus production) were plated to 70-90 % confluency at the time of transfection with complete DMEM. 18-24 hrs post-plating, cell media were replaced with DMEM supplemented with 10% FBS. For lentiviral production in each well of 6-well plate format, 1 µg of payload plasmid along with 0.2 µg pMD2.G (Addgene #12259) and 0.8 µg psPAX2 (Addgene #12260) packaging plasmid were transfected into HEK-293T cells with the Xtremegene9 transfection agent (MilliporeSigma, 06365787001) at 1 to 3 DNA to transfection agent ratio following the manufacturer’s protocol. For retroviral production in 10 cm plate, the 5 µg of payload plasmid (e.g., pMRX-IP-GFP-LC3-RFP, Addgene #84573) was transfected into Phoenix Ecotropic cells with the Xtremegene9 transfection agent at 1 to 3 DNA to transfection agent ratio following the manufacturer’s protocol. 48 hrs post-transfection, lentivirus or retrovirus supernatant was harvested, spun down at 1100 rpm for 5 min, filtered through a 0.45-µm filter, then treated to mouse KPC cells with 1:1000 dilution of 1000x polybrene transfection agent (MilliporeSigma, C788D57). 48 hrs post transduction, the transduced cell lines were selected with corresponding antibiotics: 1 mg/mL of geneticin (VWR, 100216-972) or 2 µg/mL of puromycin (Thermo Scientific, 53-79-2). As our *Brca2*-KO KPC cells already express GFP, we generated additional control and *Brca2*-KO KPC cells without GFP expression. We retrovirally introduced the autophagy reporter construct to the control and *Brca2*-KO KPC cells, then sorted the GFP+ cells 5 days post-transduction.

**Fluorescence-activated Cell Sorting (FACS)**

Five days after the retroviral transduction of the autophagy reporter construct to control and *Brca2*-KO KPC cells, GFP+ cells were sorted with the Sony SH800 Sorter. Prior to sorting, cells were dissociated into single cells, washed with 1x PBS, then resuspended into 1mL of 1x PBS with 5% BSA solution. Living, singlet, and GFP+ cells were sorted into 2 mL of FBS, then cultured in complete DMEM.

**Homologous Recombination Efficiency Assay**

The homologous recombination assay was performed according to the manufacturer’s instructions (Norgen Biotek Corporation, Catalog no. 35600). Briefly, both control and *Brca2*-deficient cells were co-transfected with dl-1 and dl-2 plasmids using X-tremeGENE9 DNA transfection reagent (Roche, Catalog no. 636779001) (Ratio transfection reagent:DNA = 6:1). After 48 hours of transfection, total genomic DNA was isolated using DNeasy Blood & Tissue Kit (Qiagen, Catalog no. 69504). To determine HR efficiency, PCR reactions were performed using supplied primer mixtures (assay and universal primers) and run under the following conditions: initial denaturation at 95°C for 3 minutes then 95°C for 15 seconds, 61°C for 15 seconds and 72°C for 15 seconds and repeated for 35 cycles then a final extension at 72°C for 5 minutes. The expected amplicon sizes for the assay and universal primer sets are 420 bp and 546 bp (to detect the backbone of dl-1 plasmid). The PCR products were then run on a 2% agarose gel and analyzed using Image J. The intensity of each recombinant product was first normalized with its internal control (dl-1 product). The HR efficiency was calculated in relative to the control cells.

**T7 Endonuclease I-based Mismatch Cleavage Assay**

To determine gene editing efficiency, genomic DNA was isolated from the cells using a lysis buffer (10 mM Tris-HCl, 50 mM KCl, 2.5 mM MgCl_2_, 0.45% Nonidet P40, 0.45% Tween 20, pH 8.3, and 10 mg/mL of ProteinaseK). PCR was performed with the isolated DNA, corresponding primers, and AmpliTaq Gold 360 Mastermix (Thermo Fisher Scientific, 4398881) following the manufacturer’s protocol. PCR products were purified with Invitrogen PureLink PCR Purification Kit (Invitrogen, K310002). For the mismatch cleavage assay (surveyor assay), two reactions were prepared per PCR product. In both reactions, the DNA was denatured and allowed to reanneal. Then, T7 endonuclease I (New England Biolabs, M0302L) was added to one set of reaction pairs and incubated for 30 min at 37˚C.

**ICE Analysis**

To verify the efficiency of FPC gene knockout for each clonally derived cell line, genomic DNA from each clone was isolated and purified with Invitrogen PureLink PCR Purification Kit (Invitrogen, K310002). For instance, to model *BRCA2* mutant PDAC patients, we lentivirally introduced Cas9 in KPC cells. A guide RNA (gRNA) targeting the mouse *Brca2* gene was cloned into LRNG vector which contains the Neo-IRES-GFP cassette, and a gRNA construct was lentivirally introduced to KPC cells (**Fig. 1A**). To ensure the loss of *BRCA2*, we clonally derived *Brca2*-KO KPC cells, then validated the presence of a frame-shift mutation in the *Brca2* gene via Sanger sequencing followed by the Inference of CRISPR Edits (ICE) analysis (5) PCR was performed to amplify the 500 bp genomic sequence flanking the gRNA target site. Amplified PCR products from each clone were sequenced with Sanger Sequencing. Synthego’s ICE CRISPR Analysis Tool was utilized to compare the control wild-type sequence to experimental sequences from each clone to determine the extent of the gene KO (5).

**CRISPR-mediated Genome Editing**

Guide RNAs were designed using CRISPR Design (http://crispr.mit.edu). gRNAs were cloned into LRG vector (Addgene #65656) (Shi et al., 2015) or lentiCRISPR v2 (Addgene #52961). LRNG vector was generated by inserting Neo- IRES cassette in front of GFP cassette in LRG vector. Lenti-Cas9- Puro (Addgene #108100) was lentivirally introduced into murine KPC cells. After antibiotics selection, gRNA was lentivirally introduced. LRNG gRNA expressing cells were selected with G418 (1 mg/mL). To make single clones, 0.5 cell per well was plated in the 96 well plate. Single cell-derived clonally expanded KPC cells were validated mutation efficiency by Surveyor assay and Sanger sequencing. In the case of the *Bub1b* gene, the highest KO score was 50 among more than 80 clones that we tested. This suggested an essential role of BUB1B as previously shown that *Bub1b* is one of the essential genes for cell survival, as a critical regulator of mitotic spindle checkpoint (6, 7). gRNA sequences used were listed in the Supplementary File 4.

**Drug Treatment Experiments**

Drugs used for this study include JQ1 (MedChemExpress, HY-13030), birabresib (MedChemExpress, HY-15743), molibresib (MedChemExpress, HY-13032), olaparib (MedChemExpress, HY10162), talazoparib (MedChemExpress, HY-16106), oxaliplatin (MedChemExpress, HY-17371), 5-FU (MedChemExpress, HY90006), gemcitabine (MedChemExpress, HY-17026), etoposide (MedChemExpress, 33419-42-0), and afatinib (MedChemExpress, HY-10261). Oxaliplatin and cisplatin were dissolved in water and other drugs were dissolved in DMSO. For EC50 analysis, cells were trypsinized to single cells. Cells were counted and diluted to 30 cells/μL in media. 50 μL of this cell media (1,500 cells per well) was plated in a 96-well plate (Thermo Fisher Scientific, FB012931). Once cells were attached (between 24 and 48 hrs after plating), drugs were added in 50 μL of media. Eight different doses plus a vehicle control were used for each drug, and three replicate wells were treated with each dose. Seventy-two hrs after the addition of the drug, cell viability was measured using AlamarBlue assay and a plate reader (Molecular Devices, SpectraMax ID5).

**CUT&RUN Assay**

For Cut and run assay, cells were trypsinized into single cells to yield 10^5^ cells for each reaction and the input sample. Enriched chromatins were extracted with reagents (Cell Signaling Technology, Cut and Run Assay Kit, 86652) according to the manufacturer’s instructions. Briefly, cell pellets were resuspended in 1mL of 1× Wash Buffer (+spermidine +protease Inhibitor cocktail). Samples were centrifuged for 3 min at 600 ×g at room temperature. For each reaction or input sample, 100 μL of 1× Wash Buffer (+spermidine +protease inhibitor cocktail) was added. 100 μL of input samples were stored at 4°C for the next DNA extraction step. 10 μL of Concanavalin A Magnetic Beads suspension were added in 100 μL Concanavalin A Bead Activation Buffer. The activated magnetic beads were placed on a magnetic rack until solution became clear and then liquid was removed. The beads were washed with Concanavalin A Bead Activation Buffer a second time and resuspended with the equal initial volume (10 μL/sample). 10 μL of activated bead suspension was added to the washed cell suspension. The samples were rotated for 5 min at room temperature. The tubes were placed on the magnetic rack then the liquid was discarded. 100 μL of Antibody Binding Buffer (+spermidine +protease inhibitor cocktail + digitonin) was added. The cell: bead suspension was aliquoted into separate 1.5 mL tubes and 2 μL of Acetyl-Histone H3 (Lys27) (D5E4) XP® Rabbit mAb (Cell Signaling Technologies, 8173) was added to each reaction. For the negative control Rabbit (DA1E) mAb IgG XP Isotype Control antibody (66362, Cut and Run Assay kit) was used. Tubes were rotated at 4°C overnight. Next day, samples were placed on the magnetic separation rack and the liquid was removed. 1 mL of Digitonin buffer (+spermatidin +digitonin +PIC) was added. Tubes were placed on the magnetic rack and the liquid was removed. 50 μL of pAG-MNase premix was added and tubes were rotated at 4°C for 1 hr. The tubes were placed on the separation magnetic rack and the liquid was discarded. 1 mL of Digitonin Buffer (+spermidine +PIC +digitonin) was added. The tubes were placed on the magnetic rack and the liquid was removed. 150 μL of Digitonin Buffer (+spermidine +PIC +digitonin) was added. pAG-MNase was activated by adding 3μL of cold Calcium Chloride and the samples were incubated at 4°C for 30 min. 150 μL of 1X Stop Buffer (+digitonin +RNAse +spike in DNA) was added and the tubes were incubated at 37°C for 10 min. The tubes were centrifuged at 4°C for 2 min at 16,000 ×g and the enriched chromatin supernatant was transferred to new tubes. For input sample preparation, 200 μl of DNA extraction buffer (+ proteinase K + RNAse A) to the 10 μL input samples. The cells were lysed by 55°C incubation for 1 hr with shaking. Lysed chromatin was then sonicated for 13 cycles (15 sec on/ 45 sec off, middle amplitude) using Bioruptor (Diagenode). After sonicated chromatin was centrifuged at 18,500g for 10 min at 4°C. DNA purification was performed with DNA Purification Buffers and Spin Columns according to the manufacturer’s instruction (Cell Signaling Technologies, 14209).

**Transfection**

siRNAs used were MISSION siRNA universal negative control #1 (Millipore Sigma, SIC001), mouse Brd2 (Millipore Sigma, SASI_Mm01_00091655), mouse Brd3 (Millipore Sigma, SASI_Mm02_00308269), mouse Brd4 (Millipore Sigma, SASI_Mm01_00116323) mouse Atg5 (Millipore Sigma, SASI_Mm01_00089196), mouse Atg12 (Millipore Sigma, SASI_Mm01_00176890) and mouse Atg16l (Millipore Sigma, SASI_Mm01_00181794). Prior to siRNA transfection, cells were seeded either on 96-well plates for cell viability assessment or 6-well plates for gene silencing verification. After 48 hrs, siRNAs targeting individual Brd2, Brd3, Brd4, Atg5, Atg12 and Atg16l were prepared in the final concentration of 20nM, while the triple siRNA combination targeting Brd2, Brd3 and Brd4 were prepared in the final concentrations of 60, 75 and 150 nM (20, 25 and 50 nM of each siRNA, respectively). All the prepared siRNAs were combined with X-tremeGENE siRNA transfection reagent (Millipore Sigma, 04476093001) according to the manufacturer’s recommendations for 72 hrs. The efficiency of siRNA knockdown was verified either by Western blot analysis or RT-qPCR.

**Quantitative Real-time PCR**

Total RNA was isolated using TRIzol reagent (Invitrogen, 15596026) and incubated for 5 min at room temperature. Chloroform was added to the samples and then thoroughly mixed them by shaking until homogeneous, followed by incubation for 5 min on ice. The samples were then centrifuged at 14,000 rpm for 15 min at 4°C. The aqueous phase was transferred to a new DNase/RNase-free tube. Isolated total RNA samples were either further processed according to the manufacturer’s guidelines or followed by column-based purification with PureLink RNA Mini Kit (Invitrogen, 12183018A) according to the manufacturer’s instructions. DNA digestion was performed on-column using PureLink DNase Set (Invitrogen, 12185010), and RNA was finally eluted with DNase/RNase-free water. The RNA concentration and purity of the samples were determined by NanoDrop ND-1000 spectrophotometer (Thermo Scientific). cDNA synthesis was performed using High-Capacity cDNA Reverse Transcription Kit (Applied Biosystems, 4368814) following the manufacturer’s guidelines. Each reaction was prepared as follows: 2 µg of RNA template, 3 µL of 10X RT buffer, 1.2 µL of 25X dNTP mix, 3 µL of 10X RT random primers, 1.5 µL of MultiScribe reverse transcriptase and 6.3 µL nuclease-free water with a total of 20 µL for 1 reaction. Reverse transcription was performed in a thermocycler as follows: 25°C for 10 min, 37°C for 120 min and 85°C for 5 min. For qPCR analysis, 1 µL of cDNA was used as template. Primers of the genes of interest (listed in the supplementary file 4), Power SYBR green master mix (Applied Biosystems, 4368702) and nuclease-free water (up to 20 µL) were added to each reaction. All qPCR reactions were performed in LightCycler 480 real-time PCR system (Roche) by setting the thermal-cycling condition as follows: 95°C for 10 min, 95°C for 15 sec and 60°C for 60 sec for 40 cycles. The relative mRNA expression levels of the genes of interested were normalized by Gapdh and calculated by the following equation: Relative mRNA expression level = 2^-ΔΔCt^.

**Immunofluorescence Staining**

Cells were cultured on an 8-well chambered cell culture slide (CELLTREAT Scientific Products, 229168). For the Rad51 foci IF experiments, doxorubicin (Spectrum Chemicals, TCI-D4193, 0.2 µM) was treated for 24 hrs. The cells were washed with PBS twice. After the final wash, the cells were fixed with 4% paraformaldehyde in PBS for 15 min at room temperature and washed with PBS once. The paraformaldehyde-fixed cells were permeabilized with 1% BSA and 0.3% Triton X-100 in PBS for 10 minutes at room temperature. The cells were then blocked with 5% BSA and 0.1% Triton X-100 in PBS for 30 minutes at room temperature. The primary antibody was prepared in PBS with 1% BSA and 0.3% Triton X-100. The antibodies used for this study include rabbit anti-LC3A/B (Cell Signaling, 12741, 1:200 dilution) and Rad51 (Cell Signaling, D4B10, 1:1000 dilution). After blocking, the cells were then incubated with primary antibody overnight at 4°C. Following 2 washes of PBS, the cells were then incubated with secondary antibody, goat anti-rabbit IgG H&L (Alexa Fluor® 594) (Abcam, ab150080, 1:1000 dilution) for 1 hr at room temperature and lastly incubated with DAPI (Genetex, GTX16206, 1:250 dilution) for 5 min at room temperature, followed by 2 washes of TBS and once with PBS. The glass slides were mounted with coverslips using Fluoromount-G slide mounting medium (Fisher Scientific, OB100-01) and imaged on a Zeiss AxioPlan II microscope using a 63X 1.4 NA objective lens together with X-Cite light source. Image acquisition was performed using a Hamamatsu ORCA-ER CCD camera and Volocity (PerkinElmer). The images were finally processed and quantified using ImageJ (NIH Image) with the Find Maxima algorithm.

**Live cell imaging and image processing**

For live-cell imaging, cells were cultured in FluoroBrite DMEM (A1896701, Gibco) supplemented with 8% FBS, 4 mM of GlutaMAX (35050061, Gibco) and 1 mM Sodium Pyruvate (11360070, Gibco). Bafilomycin A1 (S1413, Selleck Chemicals,), rapamycin (S1039, Selleck Chemicals), and JQ1 (HY-13030, MedChemExpress) were used for treating the cells. For measuring rates, 500 nM bafilomycin A1 was added along with a final concentration of 0.2 µg/mL Hoechst 33342 trihydrochloride solution (Hoechst 33342, [Invitrogen, H3570],). DMSO (472301, Sigma Aldrich) was used as a negative control for drug treatment (rapamycin or JQ1). Live cell imaging setup was previously described (8). Any position that is distorted during treatment of drugs or imaging is excluded from the analysis. An equal number of random positions were included to maintain uniform sampling for all the conditions. Images were acquired using PLAN APO LAMBDA 20X CF160 Plan Apochromat Lambda 20X objective lens, N.A. 0.75, W.D.1.0mm, F.O.V.25mm, DIC, Spring Loaded. Imaging processing steps were previously described (8). A contrast value of 16.36 was used for detecting GFP puncta while 13.25 was used for the TRITC channel. Autolysosome number was extracted by using [TRITC- TRITC HAVING (GFP AND TRITC)] binary operation. The Grow Region to intensity tool was used to account for variability in puncta size in cells in the GFP and TRITC channels. The puncta detection accuracy was confirmed through manual inspection of multiple images under various conditions. Cellpose was used for counting cells in images (9). Cellpose was implemented in Python 3.9 using a custom script and a custom-trained model was used to extract the cell count per image [https://github.com/shahlab247/image_analysis-BRCA2]. The segmentation accuracy was confirmed by manual inspection as well as by comparison with the Hoechst-based nucleus count. A custom script in Python 3.9 was used for analysis [https://github.com/shahlab247/image_analysis-BRCA2].

**Histology and Immunohistochemistry**

Harvested tumors were fixed in 10% neutral buffered formalin and replaced with 70% ethanol on the following day. Formalin-fixed paraffin-embedded (FFPE) sections with 4 µm thickness were prepared for hematoxylin and eosin (H&E) and immunohistochemistry. For immunohistochemical staining on Ki67 and cleaved caspase 3, antigen retrieval was performed for 45 min with citrate buffer at pH 6.0 in a Decloaking Chamber (Biocare Medical) at 125°C and 15 psi. Slides were blocked with normal goat serum then incubated with rabbit polyclonal Ki67 (NeoMarkers, RB1510P, 1:1000 dilution) and active cleaved caspase 3 (Promega, G748A, 1:1000 dilution) overnight at room temperature in a humidified chamber, followed by biotinylated goat anti-rabbit secondary antibody (1:1000 dilution). The Vectastain ABC kit elite kit and diaminobenzidine peroxidase substrate kit (Vector Labs) were used for amplification and visualization of signal, respectively. For TUNEL analysis, the slides were stained using TUNEL assay kit (Abcam, ab206386) according to the manufacturer’s instructions. For the quantification of IHC, three images per tumor were quantified using ImageJ software.
