## Supplementary figures for "A new vulnerability to BET inhibition due to enhanced autophagy in BRCA2 deficient pancreatic cancer"

#### Supplementary Figure 1

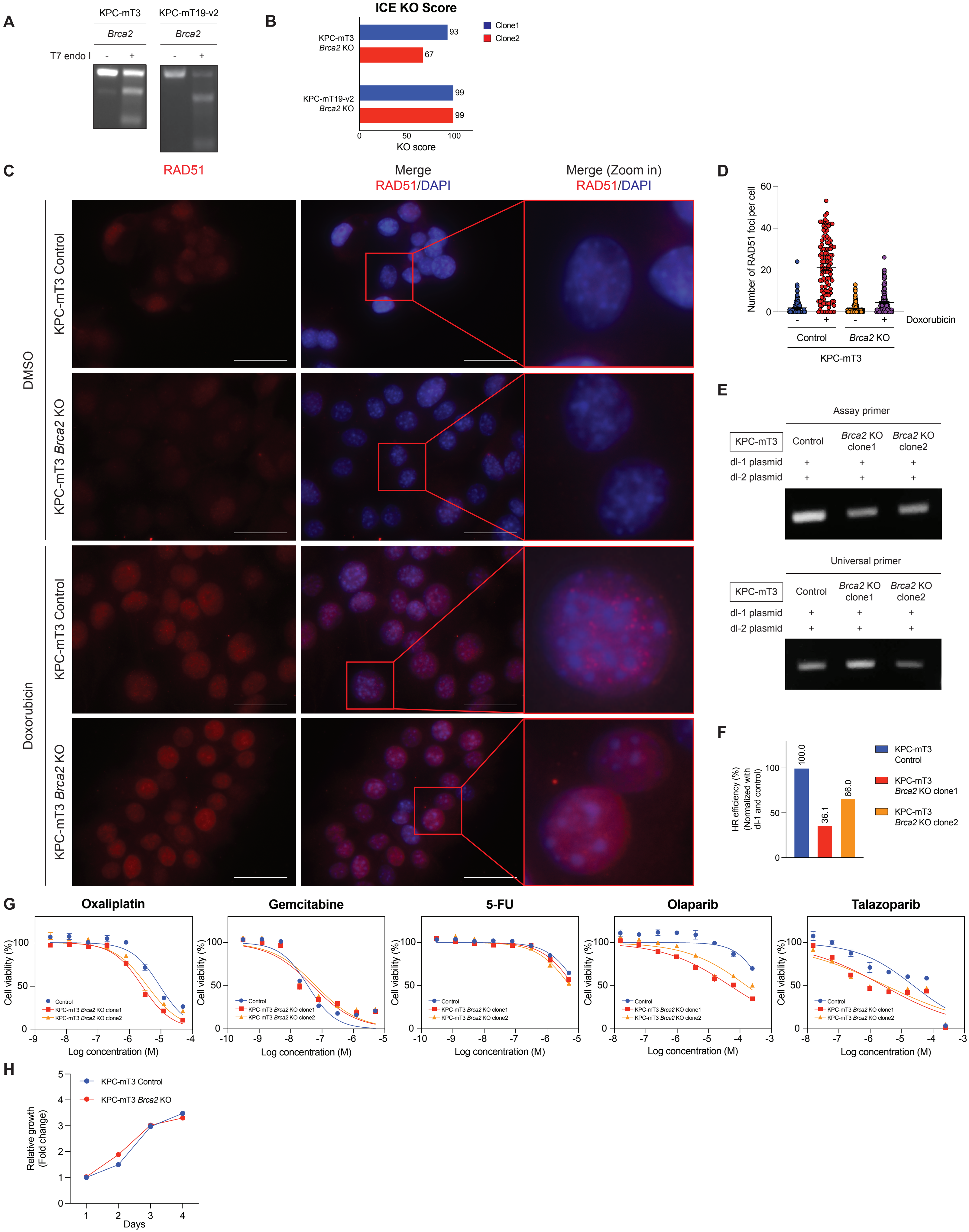

### Supplementary Figure 2

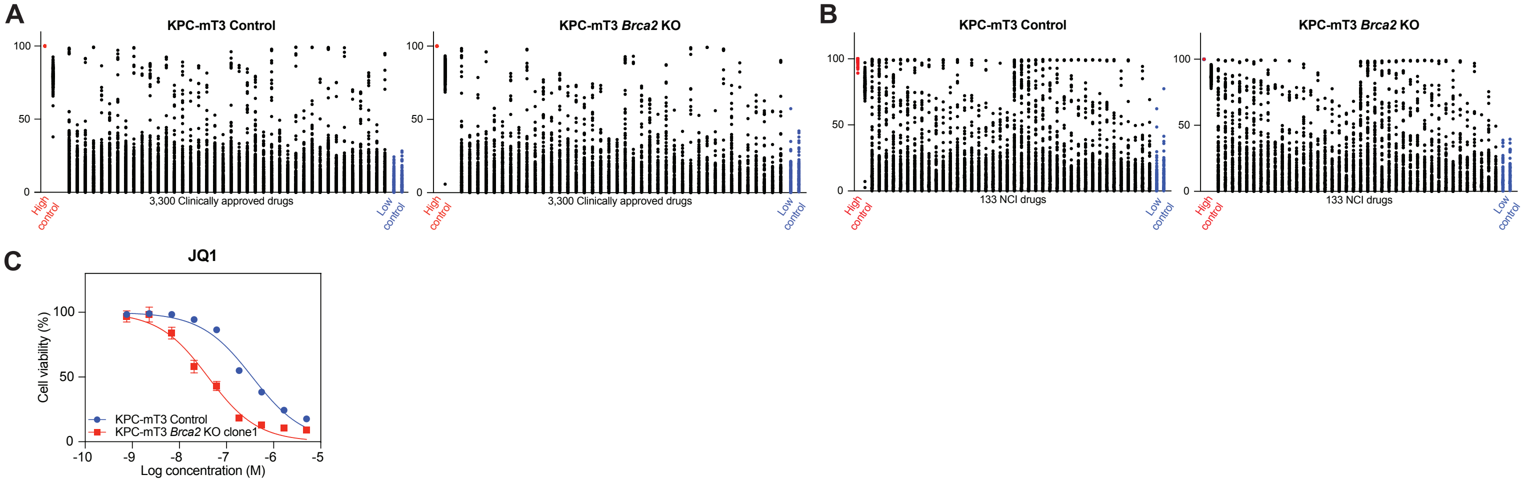


| | Control | Brca2 KO |
|---|---|---|
| **HillSlope** | -1.223 | -0.9907 |
| **IC50** | 1.880e-007 | 2.862e-008 |

### Supplementary Figure 3

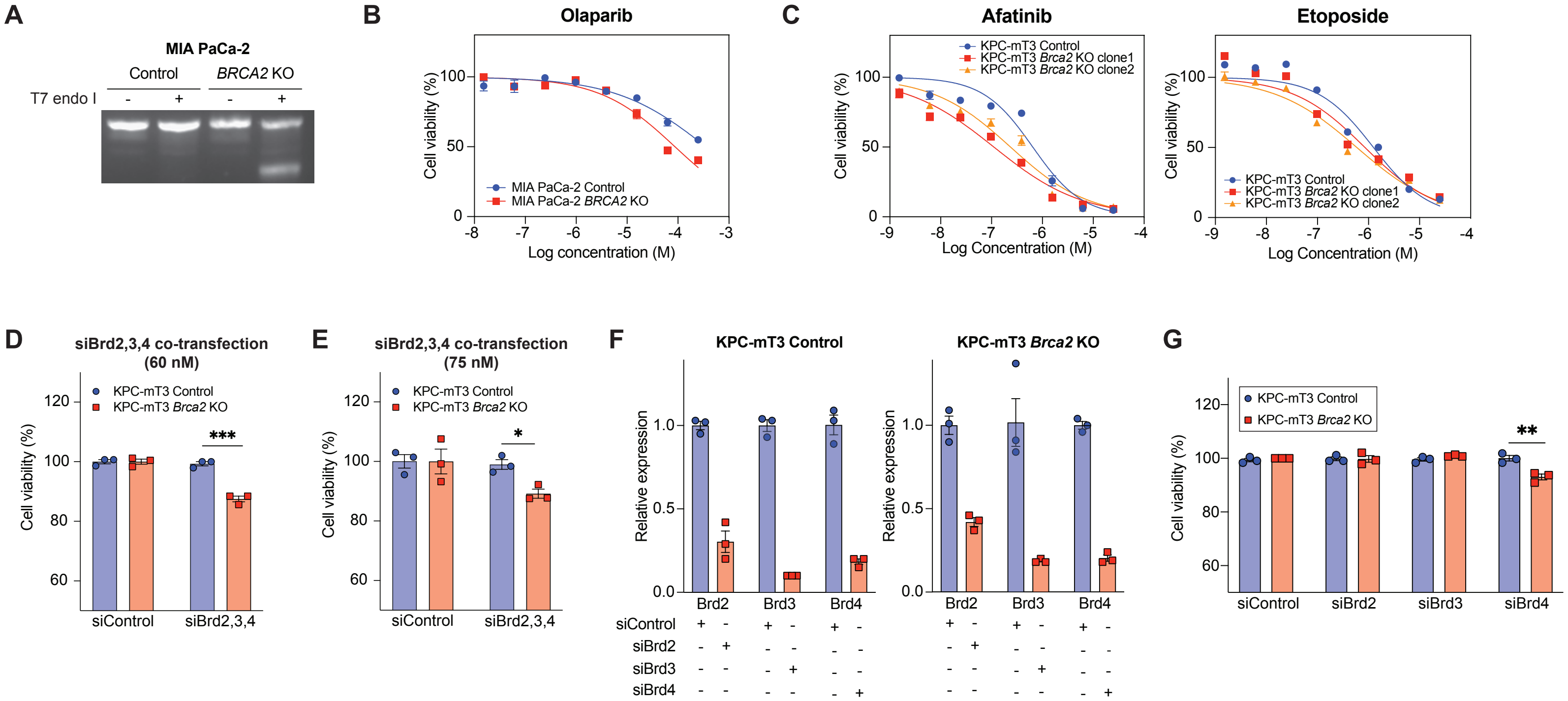

#### Supplementary Figure 4

A

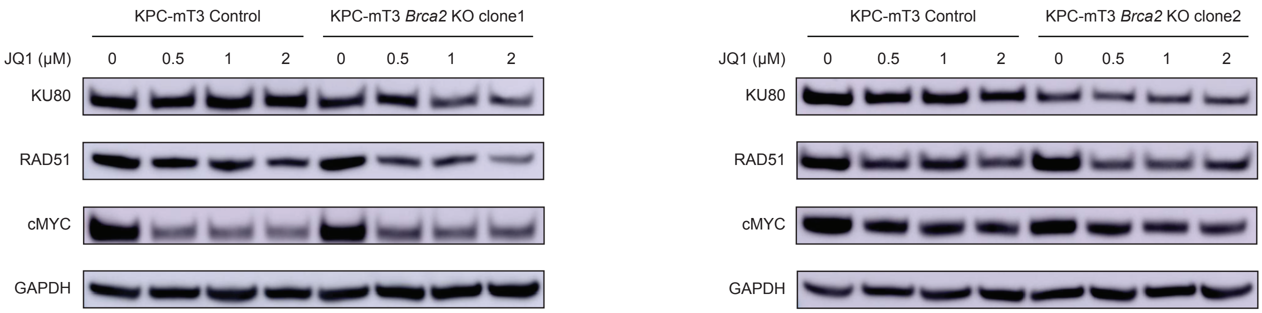


B

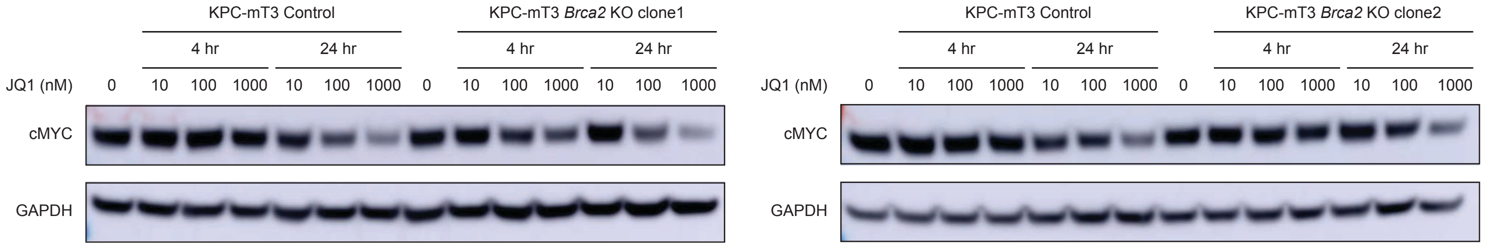

**Supplementary Figure 5**

**A**

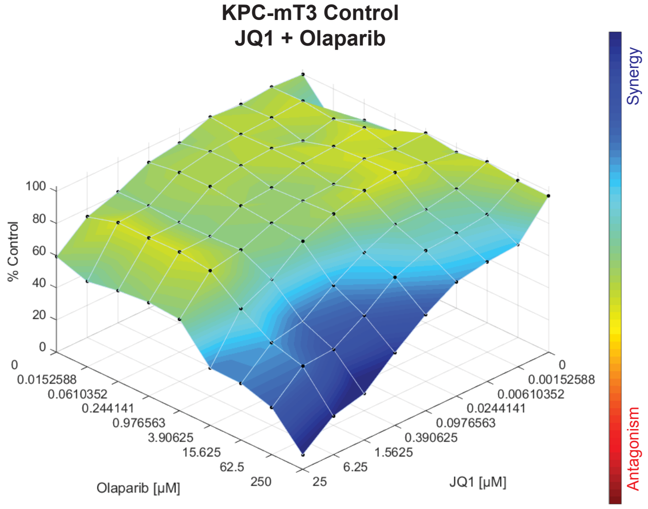


**B**

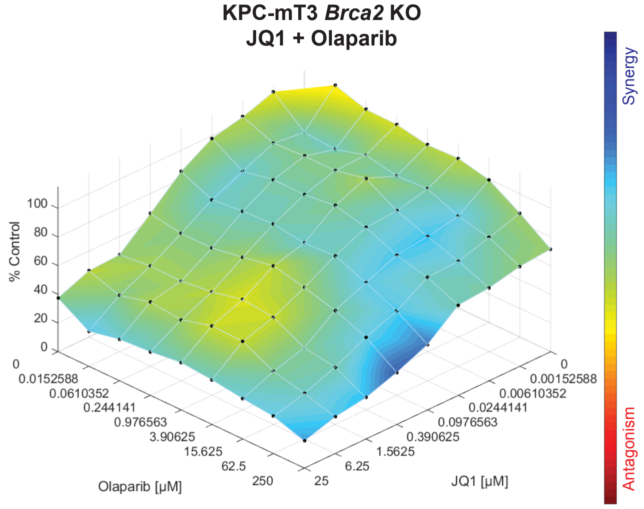

### Supplementary Figure 6

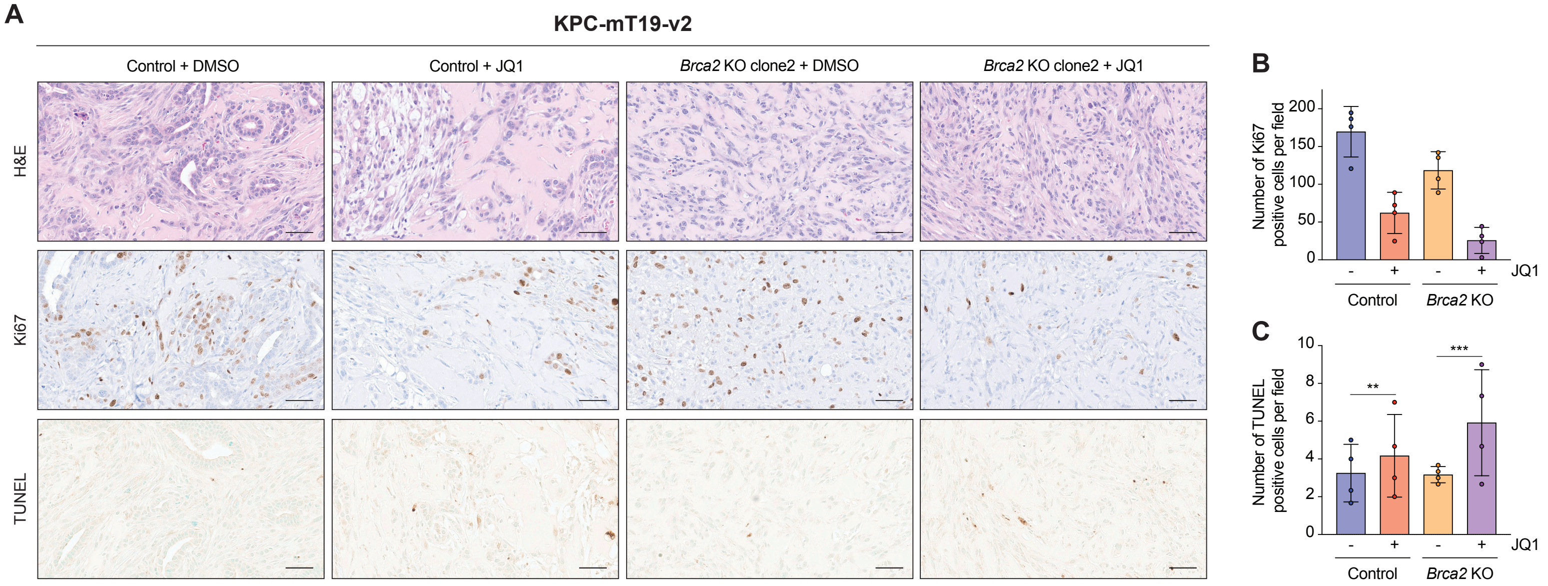


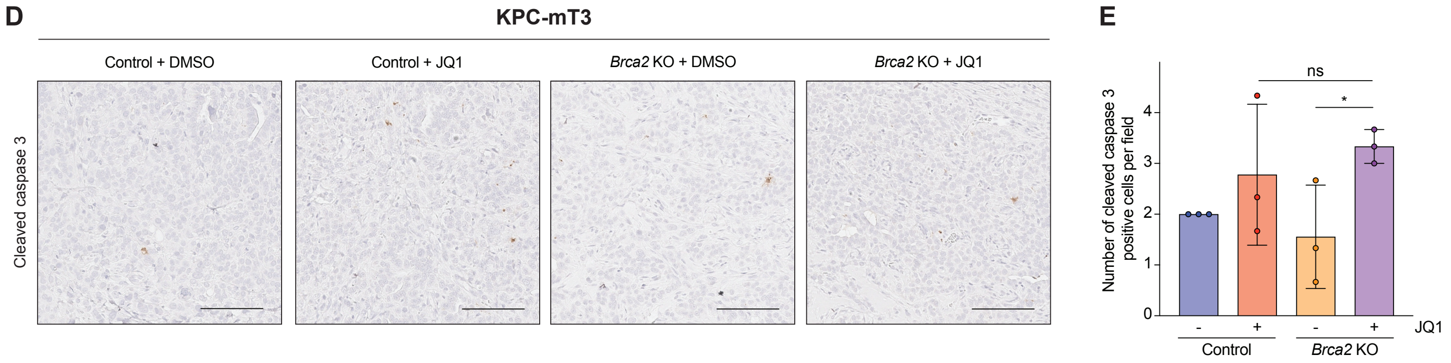

#### Supplementary Figure 7

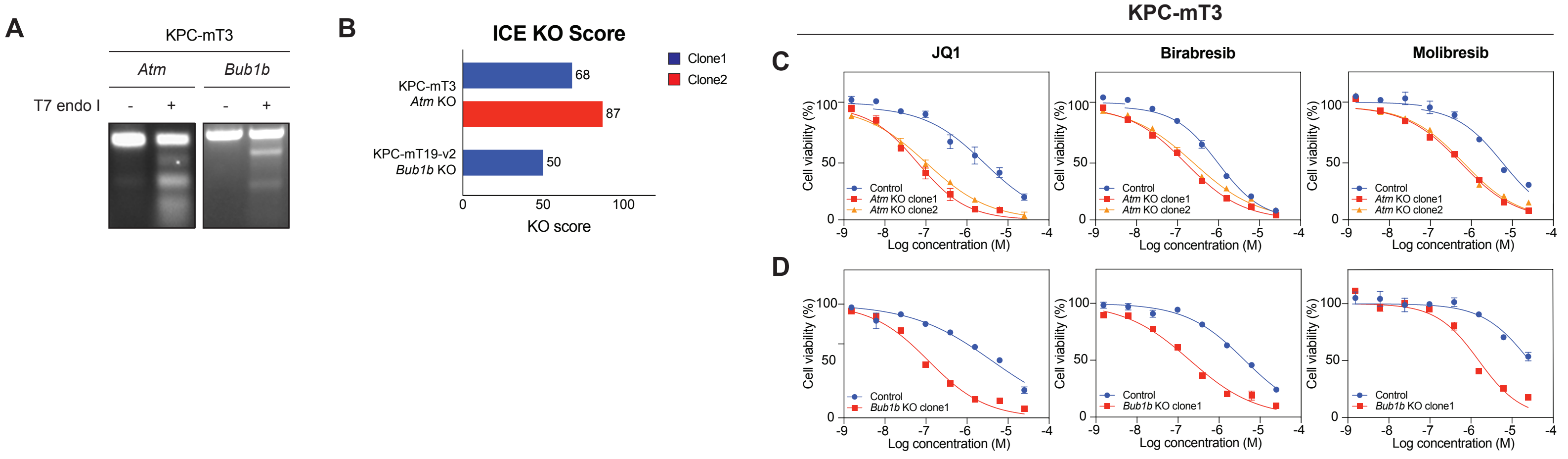

**Supplementary Figure 8**

A

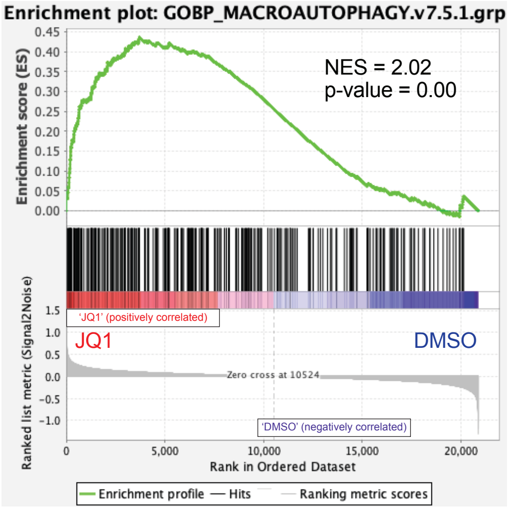


B

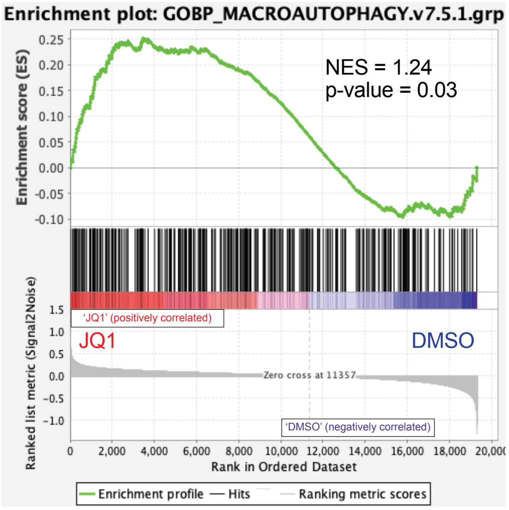

**Supplementary Figure 9**

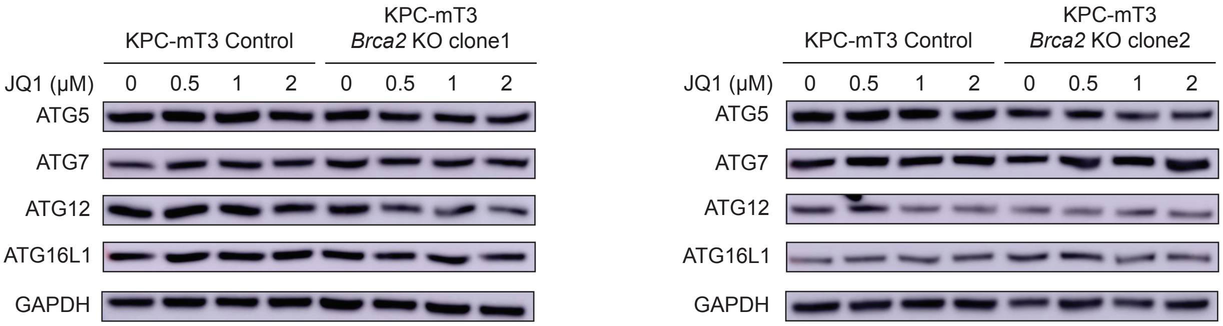

#### Supplementary Figure 10

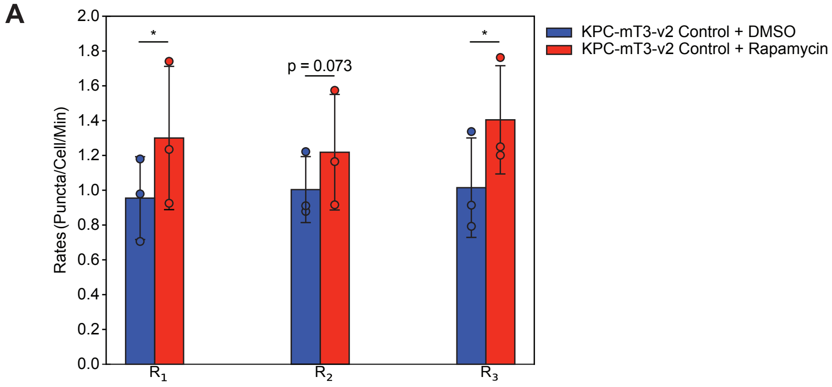


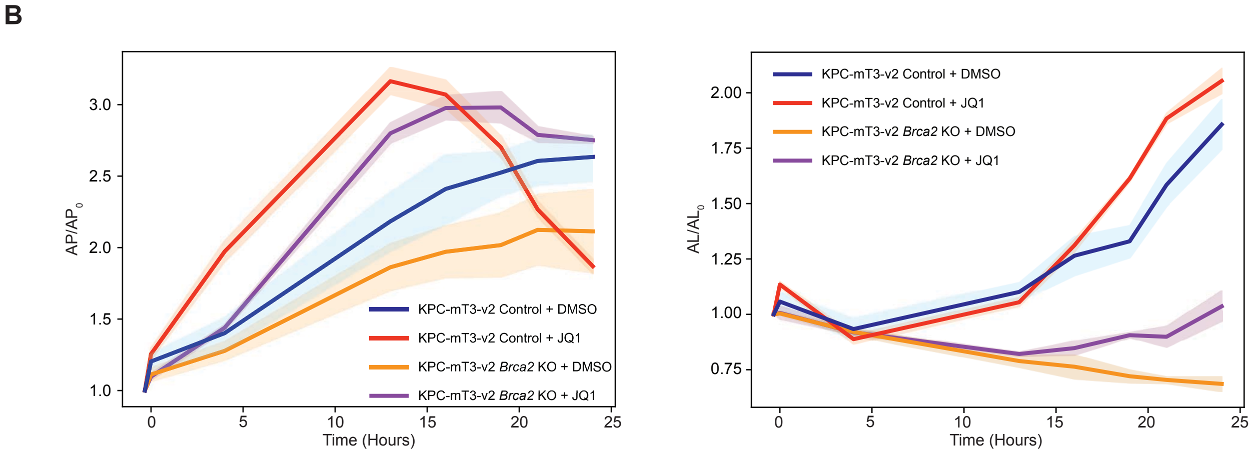

**Supplementary Figure 11**

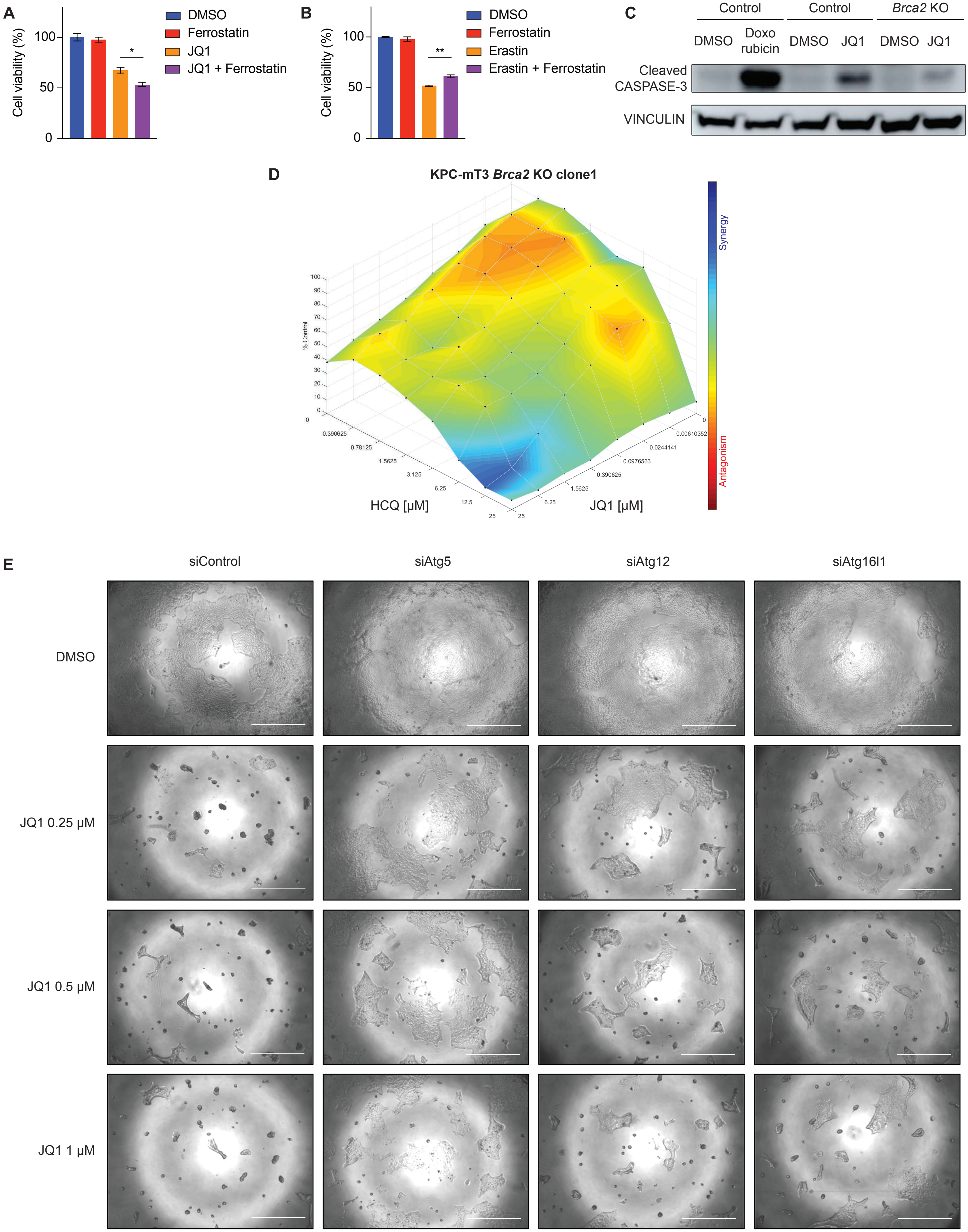
